# A consensus motif in *ASH1* and further transcripts unifies several RNA motifs required for interaction with the She2p/She3p transport machinery and mRNA localization in yeast

**DOI:** 10.1101/2024.03.10.584336

**Authors:** Markus Seiler, Annika Niedner, Simone Heber, Michael Feldbrügge, Ralf-Peter Jansen, Dierk Niessing, Kathi Zarnack

**Author notes:** Equal contribution (Markus Seiler, Annika Niedner).

## Abstract

Asymmetric localization of the *ASH1* transcript is a central step in the regulation of mating type switching in *Saccharomyces cerevisae* and a paradigm for localized mRNAs specifically recognized by the She2p/3p transport machinery. Four RNA elements in *ASH1* are known to mediate bud localization, but it remained unclear what a consensus motif of all four She2p/3p recognition sites (SRS) might look like. In this study, we derive the SRS consensus motif, which is characterized by an L-shaped, double-stranded RNA conformation with two cytosine-containing sequence motifs at conserved positions and a central asymmetric bulge. It is noteworthy that the spatial arrangement of these features can be retained, despite their altered order within the nucleotide sequence. We termed these variations “configurations” and confirmed in biochemical studies that SRS function in *ASH1* elements is preserved when changing configurations. We tested other known and predicted SRS elements in target transcripts of She2p/3p by biochemical and biophysical studies and confirmed the conserved SRS function even in switched configuration. Consistently, we present the *SRS instance collection*, which includes further predictions of SRS instances of transcript targets of the She2p/3p translocation pathway.

## Introduction

The active intracellular transport of effector molecules contributes to the high degree of organization in life and asymmetric mRNA distribution is one way to ensure correct spatiotemporal expression of encoded proteins^1^. In recent years, it has been studied in various polarized cells, in *D. melanogaster* and *Xenopus laevis* oocytes and embryos^2–4^, and also in mammalian fibroblasts^5^ and neurons^6^. mRNA transport is involved in embryonic patterning, cellular polarity, and asymmetric cell division. Failure of mRNAs to be properly transferred has been shown to cause cellular dysfunction and disease^7^. In these pathways, mRNAs are recognized by the cellular translocation machinery via *cis*-acting elements in their sequences, called zip-codes or localization elements that may be determined by short RNA sequence motifs, conformational folds^8–10^ and spatial restraints between these features^11^. These zip-codes are recognized by RNA-binding proteins that target the mRNAs to their final destination^12^. Within the mRNAs, the zip-codes often reside in the 3’ untranslated regions (3’ UTRs) of mRNAs, but also in 5’ UTRs and coding regions^13–18^.

Budding yeast (*S. cerevisiae*) has evolved as a model organism for mechanistic analyses on specific targeting of mRNAs to the bud of emerging daughter cells^19,20^. The current model of how the RNA transport machinery is built suggests a step-wise assembly of its core compounds: She2p interacts specifically with its target mRNA and recruits the adapter protein She3p together with the type V myosin Myo4p^21–24^. This mRNA ribonucleoprotein (mRNP) complex is then transported along the actin cytoskeleton. The paradigm, the yeast *ASH1* mRNA, is localized to the bud tip enabling the asymmetric distribution of its encoded transcription factor Ash1p into the daughter cell nucleus where it represses mating-type switching^25–28^. To date, over twenty transcripts have been reported to be targeted via the She2p/3p machinery. Besides the well-characterized She2p/3p target mRNAs *ASH1*, *EAR1*, and *WSC2*, further candidates were identified using combined RNA immunoprecipitation and microarray techniques^24,29^. In some target mRNAs, the regions containing putative zip-codes had been narrowed down by yeast three-hybrid and *in vivo* localization studies^30^. However, besides of *ASH1*, precise positions of the She2p/3p recognition sites (SRS) remained unknown in most cases. Within the *ASH1* mRNA, four localization elements with intrinsic SRS function were mapped in accurate fashion^13,14,19^. Three of these (E1, E2A and E2B) were shown to reside within the coding region, whereas E3 overlapped the stop codon and extended into the 3’ UTR (**Supplementary Fig. 1A**, **Tab. 1**). Each *ASH1* SRS element was sufficient for triggering the assembly of the She2p/3p complex and for localizing a reporter RNA to the bud and was predicted to exhibit RNA folds with extended stem-loop structures. An *in-vivo* screen with mutated *ASH1* fragments supported the notion that a three-dimensional fold was involved in zip-code function. The fold was described as a stem with conserved nucleotides on either side: a singular cytosine and a CGA triplet, with a distance of six nucleotides between the cytosines. Searching for similar RNA folds identified functional SRS elements in *IST2* and *EAR1*^10^. However, these folds lacked an apparent consistency with some of the four *ASH1* SRS elements and a coherent model for specific SRS recognition by the She2p/3p transport machinery has remained elusive. Recently, the interaction interface between E3 and She2p/3p was resolved using X-ray crystallography: a symmetric tetramer of She2p exhibited two interfaces interacting with E3 on opposite sides of the complex^8^. As suggested before^31^, the affinity was increased by She3p in a synergistic manner. Intriguingly, the ternary complex formation was accompanied by an induced fit in the SRS element: For the unbound E3 RNA, SHAPE experiments suggested an elongated conformation^8^. When interacting with She2p/3p, E3 displayed a sharp turn in one of its bulge regions, resulting in a kinked global conformation that was also confirmed by NMR structures.

**TABLE 1.** Overview on the four localization elements of *ASH1*. Positions and configuration classification are listed according to the SRS consensus motif

| SRS name | SRS config I | SRS config II | SRS AA motif in Lbulge | SRS start | SRS end |
| --- | --- | --- | --- | --- | --- |
| E2 |  | II | yes | 631 | 686 |
| E2A |  | II | yes | 1118 | 1178 |
| E2B |  | II | yes | 1275 | 1312 |
| E3 | I |  | yes | 1774 | 1817 |

In this study, we define a coherent SRS consensus motif that unifies the features of E1, E2A, E2B and E3 in *ASH1* and that also classifies SRS elements of further bud-localizing transcripts (*WSC2*, *EAR1, IST2*). It exhibits a kinked stem-asymmetric bulge-stem-bulge structure, with the recurrent singular cytosine in the asymmetric bulge and the CGA motif in a second bulge. One characteristic of the consensus motif is its arrangement of features, conserved in the overall spatial arrangement but present in two variations (“configurations”) in linear sequences. Electrophoretic mobility shift assays and surface plasmon resonance studies showed that E3 and other SRS elements could be switched between configurations without impairing She2/3p binding. Using computational screening, we identified potential SRS instances in further twelve transcripts known to interact with She2/3p. Finally, we introduce the *SRS instance collection*, describing in detail predicted SRS elements and their positions in further RNA targets of the She2p/3p translocation pathway.

## Results

### Bioinformatics predictions reveal a consensus capturing all She2/3p recognition sites of *ASH1*

We wondered if a general concept may exist to describe all four localization elements of the *ASH1* mRNA that act as She2p/3p recognition sites (SRS). For this study, we could rely on a recently published crystal structure that showed in detail how the *ASH1* localization element E3 interacts in a kinked stem structure with She2p/3p^8^. Of note, this interaction involved a molecularly induced fit upon binding that resulted in an asymmetric bulge in the RNA stem with one of its functional cytosines deeply buried inside the protein interface. The asymmetric bulge of E3 displayed a large stretch (“Lbulge”) and a small one (“Sbulge”), with a length ratio of 6 nt: 3 nt (**Fig. 1A, right**) or 4 nt: 1 nt, respectively, if considering atypical base pairing inside the bulge as further base pairs. We hypothesized that the ability to form such an asymmetric bulge could be a pre-requisite of the E3 conformation, whose composition would require different lengths to be kinked. In a next step, we performed a systematic survey of such arrangements in further SRS elements of *ASH1*. However, common RNA conformation predictors do not consider precise structural rearrangements due to RNA-protein interaction very well. In a step-wise strategy (**Supplementary Fig. 2, Material and Methods),** we first evaluated which of the commonly used prediction tools would predict the best approximation of the kinked E3 conformation in complex with She2p/She3p^8^ just from the single E3 nucleotide sequence. Of note, we found *Kinefold*^32^ to predict the asymmetric bulge in close approximation (5 nt: 2 nt) that could even be closer approximated when considering atypical base pairing as found in the asymmetric bulge of the crystal structure, resulting in (6 nt: 3 nt) (**Fig.1**, **Supplementary Fig. 3A**).

**Figure 1.**
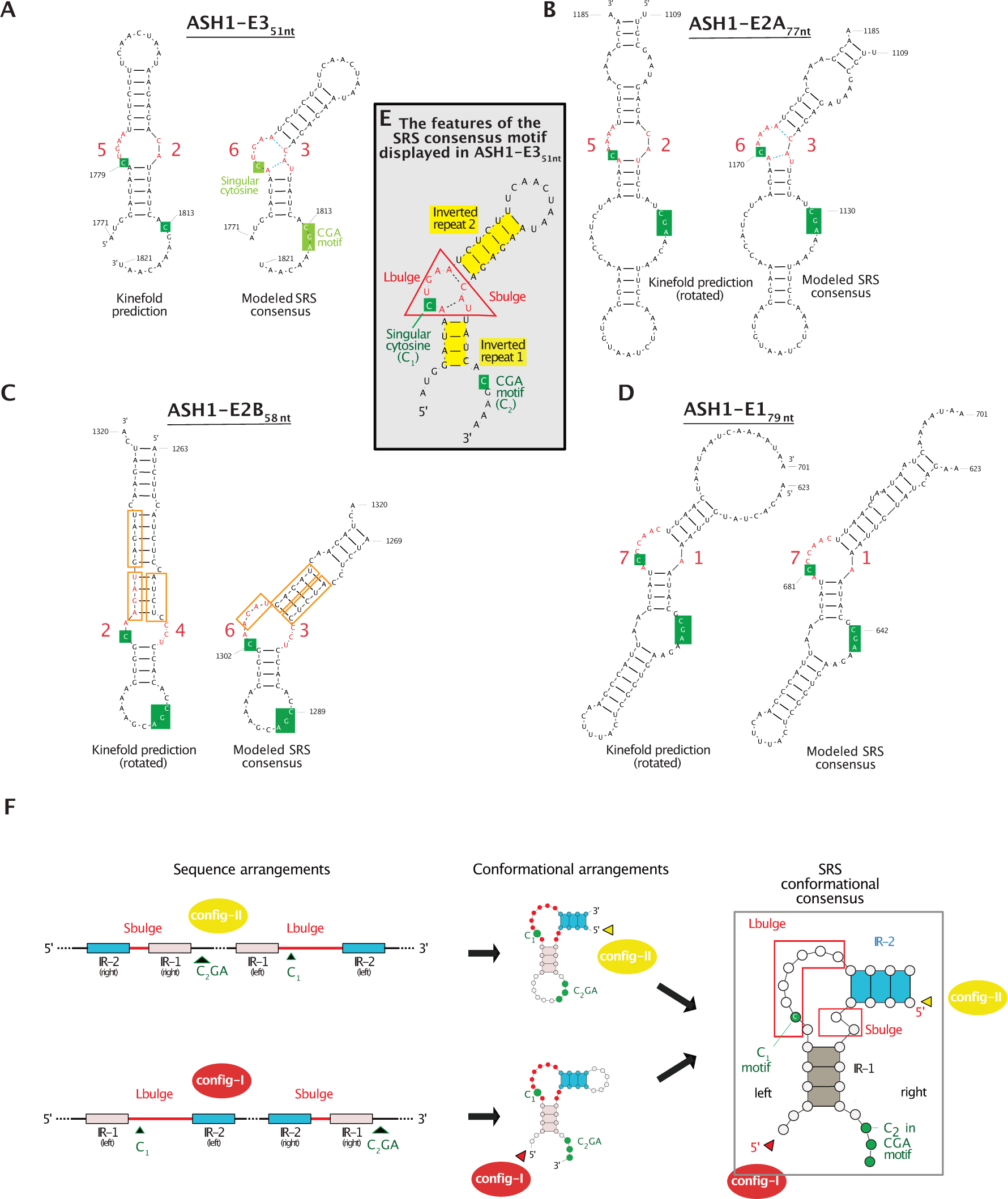
The SRS consensus model describes all four *ASH1* SRS elements. **A** Left, conformational prediction of E3_1771-1821_ by Kinefold displays the asymmetric bulge as a strand of 5 nt (‘Lbulge’) juxtaposed with a shorter strand (‘Sbulge’) both indicated in red, previously known functional motifs in dark green. Right, the conformation of E3 referred to its crystal structure bound to She2p/3p (**PDB: 5NEW**) displayed a rearranged the asymmetric bulge with atypical base-pairing, with a length ratio of 6 nt: 3 nt (with atypical base-pairs dashed in blue, but ignored in the count), and with further positions of sequence motifs adapted to analogous functional residues as studied in E2A, E2B and E3 (indicated in light green). **B** Left, Kinefold prediction and 180° rotation of E2A_1109-1185_ revealed an asymmetric bulge with the same 5 nt: 2 nt ratio as in E3. Right, the high degree of sequence similarity (**Supplementary Fig. 1**) enabled a modeling in tight accordance to the E3 conformation of the crystal structure that even comprised the uncommon base-pairing C_1122_-C_1174_ and the A_1123_-A_1169_ base-stacking (dashed in blue). **C** Left, the optimal conformation of E2B_1263-1320_ after 180 ° rotation showed internal repeats (left, boxed in orange) that could be re-modeled according to Kinefold predictions (right) but with increased energy content (−10.3 kcal/mol vs.-7.4 kcal/mol). **D** SRS element E1_623-701_ after Kinefold prediction and 180° rotation (left) and with adapted conformation (right). **E** Scheme of the features of all four modeled SRS elements, projected on E3_1771-1821_. An asymmetric bulge (red) is framed by two inverted repeats (yellow), with sequence motifs indicated in green. **F** Schematic presentation of the SRS consensus motif with its features displayed in both configurations (IR1 boxed in grey, IR2 boxed in blue, Lbulge and Sbulge in red, sequence motifs indicated by green labeling). Regarding the two configurations, their different order of features (left) results in a overall spatial arrangement that is conserved but with varying positioning of 5’ start and 3’ end in its conformations (middle). Right, conformational view on the unified model of the SRS consensus motif.

For E2A, we found the predicted conformation with an asymmetric bulge with exactly the same length ratio as found in E3 (5 nt: 2 nt) using *Kinefold* (**Fig. 1B left**). Of note, after 180° rotation, the predicted E2A conformation harmonized perfectly with the E3 conformation (**Fig. 1A right, 1B right**). Furthermore, this 180° rotation caused the previously diverging sequence motifs of E3 to fit now to the single cytosine and CGA motifs of E2A. Surprisingly, sequence comparison revealed 89% sequence identity between E3 and E2A in the asymmetric bulge and flanking regions (**Supplementary Fig. 1B**). For E1, the 180° rotation of its predicted conformation resulted in an arrangement comparable with E3 (**Fig. 1D**), but with different length proportions of the asymmetric bulge and a lower degree of sequence identity. These findings suggested one blueprint, with a common overall spatial arrangement of its (conformational and sequence) features in E3, E2A and E1 that we referred to as “SRS consensus motif” (**Fig. 1E,F**). Of note, despite of this conserved spatial arrangement of features, the linear order of features was interchangeable in the nucleotide sequences (**Fig. 1F**). This was due to the fact that the conformations of E3, E2A and E1 are harmonized by a 180 degree rotational step of E3. Subsequently, the 5’ start of E3 was at the opposite site of the SRS consensus motif than for E2A and E1, resulting in different feature orders in the nucleotide sequence (**Fig. 1E**). We termed these variations “configurations”, with “config-I” observed in E3 and “config-II” in E2A and E1. Further, we could harmonize the annotation of the sequence motifs in E3 by labeling the single cytosine motif in the asymmetric bulge as C_1_ and the cytosine in the CGA motif as C_2_ (**Fig. 1A, right, 1E**), analogously to E2A and E1. For E2B, at a first glance, the conformational prediction of E2B displayed the hypothesized “Lbulge” even shorter than the “Sbulge” (**Fig. 1C, left**). Remarkably, Kinefold predicted an alternative conformation with an asymmetric bulge in agreement with the SRS consensus motif (6 nt: 3 nt), but with a higher content of energy (calculated free energy = −10.3 kcal/mol versus −7.4 kcal/mol), which could be rearranged by sliding between sequence repeats in one inverted repeat (**Fig. 1C, right**). Altogether, we had derived a unified model of the SRS consensus motif (see **Info**-**Box**). In a subsequent step, we could confirm that the model was sufficient to summarize previous functional studies with mutationally-derived decreases in RNA localization in all four *ASH1* SRS elements^10,13,14,25^ (**Material and Methods)**. For an overview, the distinct modeling steps (initially characterized elements, Kinefold prediction, rotation, and mutual feature alignments) are displayed in (**Fig. 2**).

**Figure 2.**
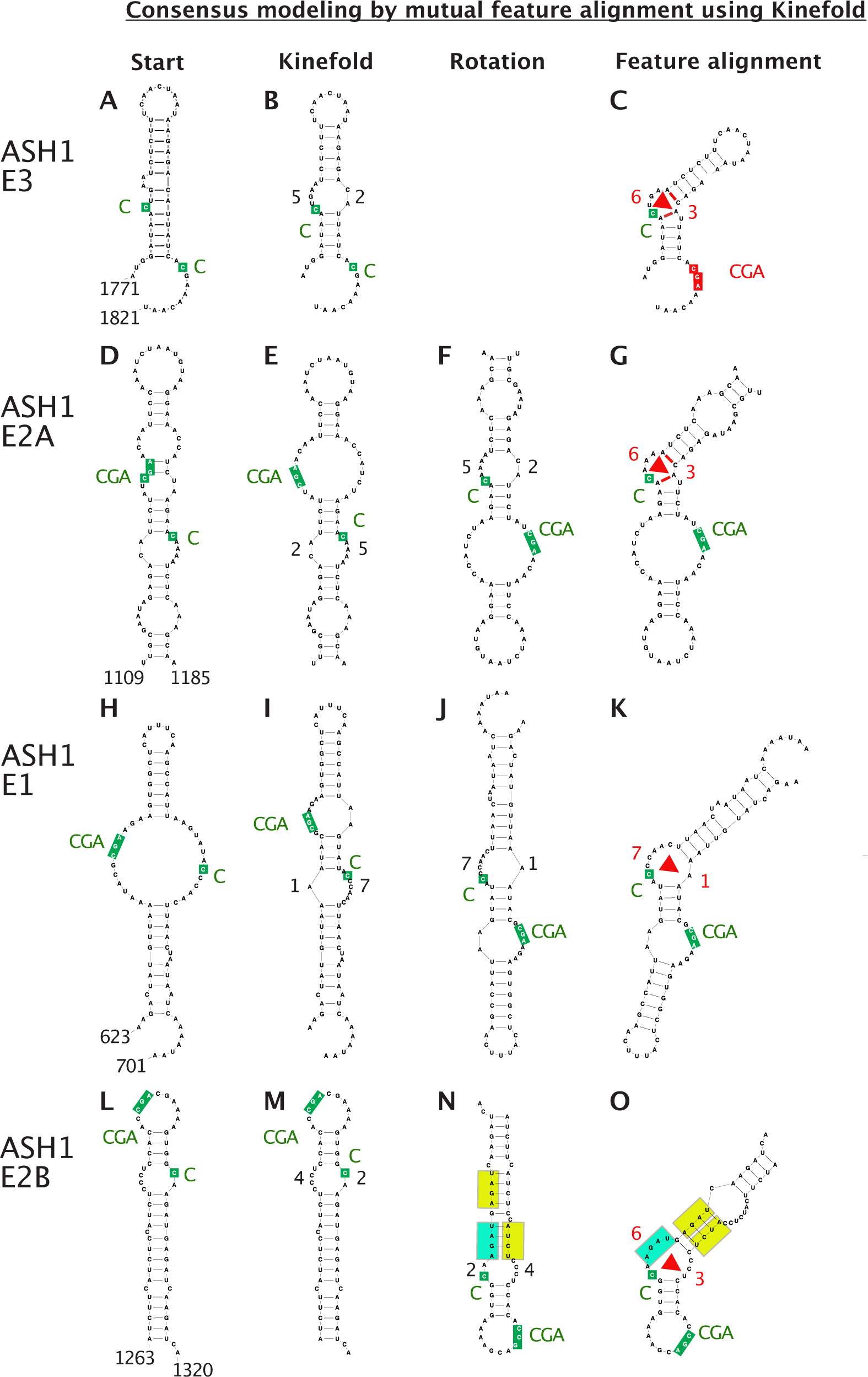
The modeling of the SRS consensus involves choice of appropriate conformation prediction tool, rotation and mutual feature alignment of the four SRS elements in *ASH1.* The overview demonstrates the effects of the three steps of modeling on the extended sizes of the *ASH1* SRS elements as they were characterized before. Whereas the initially known conformations displayed a huge degree of heterogeneity (**A, D, H, L**), the conformation prediction tool Kinefold tolerated asymmetric bulges in each SRS element (**B, E, I, M**). For E3, using the published crystal structure as a blue print, its conformational prediction (**B**) was adapted, with altered conformation of the asymmetric bulge (resulting in atypical base pairings, marked in red, **C**). Its final asymmetric feature is indicated by a red triangle, with the numbers of nucleotides displayed for both, Lbulge and Sbulge. By later analyses, we could assign one of its functional cytosines as being analogous to already known CGA sequence motifs with other SRS elements (indicated in red). For E2A, E1 and E2B, it was the rotation of the initial predictions that harmonized the respective conformations with E3 (**F, J, N**). For E2A, due to its remarkable sequence identity with E3, we could also introduce the same atypical base pairings (marked in red, **G**). As mentioned before, E2B represented an outlier: it was a alternative conformation prediction that could be harmonized with E3, suggesting the need of a further intramolecular rearrangement to align the inverted boxes (marked in yellow, **N**, **O**).

### The two configurations are interchangeable in E3 without loss-of-function *in-vitro*

One unexpected characteristic of the SRS consensus motif was the spatial conservation of its features in its overall conformation while the order of features in the nucleotide sequences was found to be swapped for the different configurations. To validate if we had grasped the essence of the consensus, we investigated if an artificial switch between the different sequence orders (and the configurations) would conserve SRS function. For this purpose, we used the well-characterized construct E3_51nt_wt^8^ and switched its feature order in the nucleotide sequence (config-I) to design a config-II construct (E3_51nt_switch, **Fig. 3A**). Based on the observation that the E3 crystal structure exhibited an interaction of She2p/3p with C_2_G but not with C_2_GA, and for comparability reasons with previous studies, we limited the construct design to C_2_G. Using an electrophoretic mobility shift assay (EMSA), we examined E3_51nt_switch for interaction with She2p and She3p in a titration approach. We used the fusion protein She2p-She3p, which comprised both, She2p and a She3p fragment and was demonstrated to be sufficient to mediate the synergistic binding of both proteins^8^. For the wildtype construct E3_51nt_wt, we observed an interaction strength at the expected nanomolar protein concentration (**Fig. 3A**)^8^. As a control, we tested the construct E3_51nt_M6 characterized by reduced interaction^8^ due to a C_1813_G substitution of C_2_. In agreement of our model, E3_51nt_wt and E3_51nt_switch, interacted with She2p-She3p at comparable nanomolar protein concentration (**Fig. 3A**). These similar binding affinities contrasted with E3_51nt_M6, which displayed severely impaired interaction.

**Figure 3.**
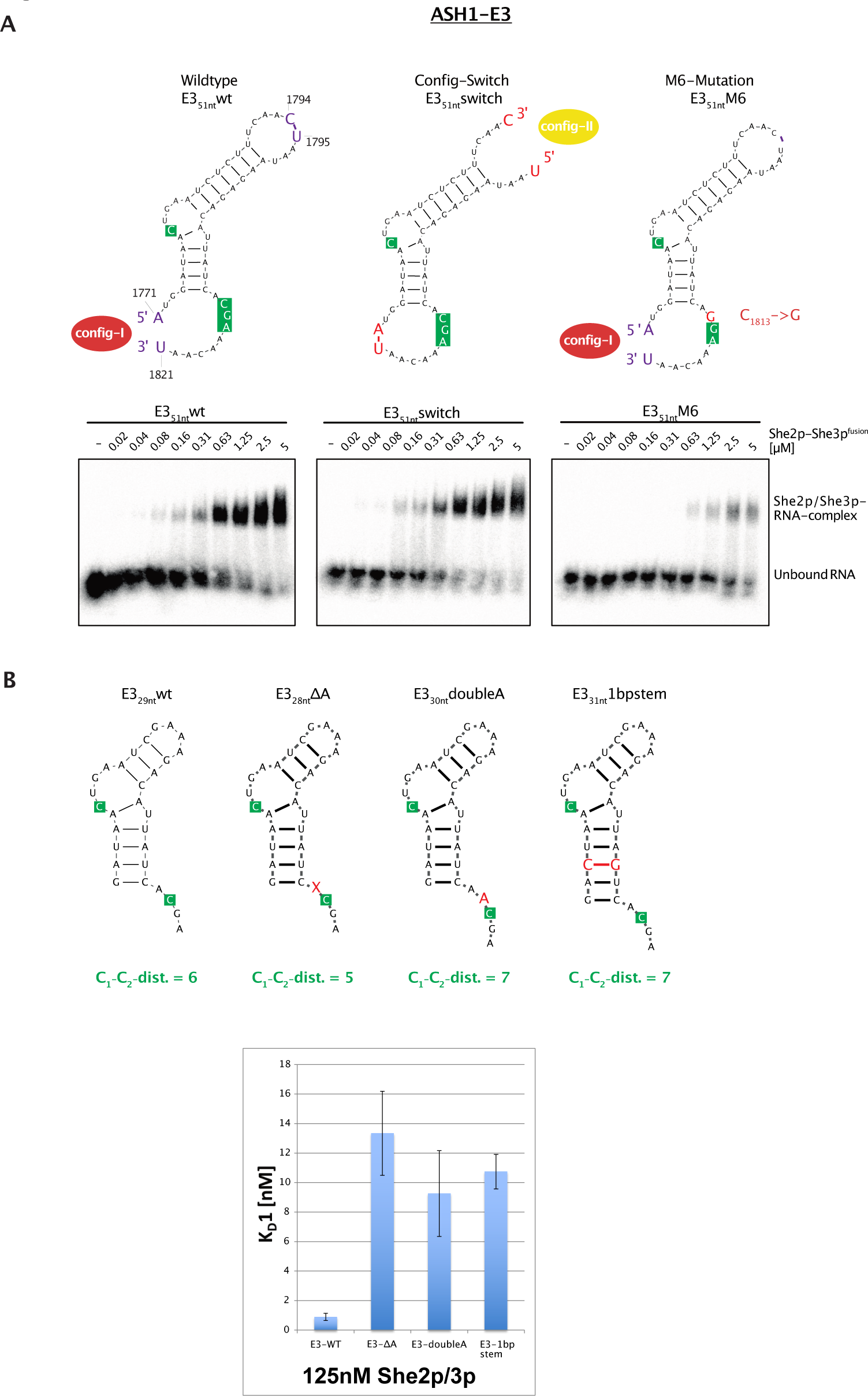
The configuration switch in E3 displays conserved function and surface plasmon resonance analysis confirms the optimal C_1_-C_2_ distance of 6 nt in minimal constructs of E3. **A** Representative results of electrophoretic mobility shift assays (EMSA) of triplicates to analyze complex formation of wild-type and variant constructs of radioactive ASH1 E3_51nt_ RNA with the She2p-She3p fusion protein (She2p_6-246_-(GGSGG)_2_-She3p_331-405_) in increasing concentration (0.02 - 5 μM): Wildtype E3_1771-1821_ (left), construct with switch of E3_1771-1821_ from config-I to config-II (middle) and control M6 mutational variant of E3_1771-1821_ with decreased SRS function^8^. **B** Results of surface plasmon resonance analysis (SRP), performed in triplicates to analyze the minimal E3 constructs and alterations in their C_1_-C_2_ distance. The minimal construct E3_28nt_wt was re-designed to create constructs with differing C1-C2 distances (5 and 7 nt) and was incubated with She2p-She3p fusion protein (125 nM).

Since the E3_51nt_M6 mutant showed impaired but still considerable binding to She2p-She3p-fusion protein, we used the full-length proteins to assess the binding features of E3 elements. Incubation with constant She3p concentration of 0.56 μM and titration of She2p (0.02 - 5 μM) confirmed that both constructs, E3_51nt_wt and E3_51nt_switch, supported binding even at the lowest She2p concentration used in this experiment (0.02 µM). In contrast, binding of the full-length proteins to E3_51nt_M6 was abolished at concentrations up to 5 µM (**Supplementary Fig. 4B**). The same was true for a reciprocal approach, in which increasing concentrations of She3p (0.02 - 5 μM) were tested in the presence of high amounts of She2p (5 μM) (**Supplementary Fig. 4C**). Moreover, both experiments showed that She3p efficiently binds E3 in the absence of She2p. Surprisingly, this binding was sensitive to mutation of C_2_. In conclusion, these studies successfully confirmed that E3 can be switched from config-I to config-II without compromising its functionality to interact with She2p/3p.

When we analogously examined E2A and E1, we reversed the direction of switching (now from config-II to config-I), resulting in the constructs E1_27nt_switch and E2A_32nt_switch (**Supplementary Fig. 5**). Again, in EMSA validations, we could demonstrate that both constructs largely exhibited the binding properties to She2p-She3p-fusion. In summary, the “config-switch” approach supported our model, which suggested that SRS features could be rearranged in the RNA sequence while retaining function. We could demonstrate that E1, E2A, and E3 recognized She2p/3p regardless of configuration, demonstrating the essential characteristics of RNA-She2p/3p recognition were reflected in the SRS consensus motif. As a conclusion, these findings indicate that SRS function requires very few conserved bases in defined positions of a stem-loop, which must accommodate a bulge that is flexible enough to open up upon binding by She2p-She3p to adopt a kinked L-shaped conformation in the co-complex.

In a fine-grained approach, we used the analytical power of surface plasmon resonance (SPR), to investigate the influence of the functional cytosines C_1_ and C_2_ on SRS function. Previous studies had determined a preferred distance of six nucleotides between these cytosines^10^. Using the functionally minimal^8^ construct E3_29nt_, we introduced varying distances between C_1_ and C_2_ and performed titration series in a surface plasmon resonance experiment with a She2p-She3p-fusion construct (125 nM) (**Fig. 3B**). As a result, for the wildtype construct E3_29nt_wt, we observed a binding affinity of approx. K_D_ = 0.89 nM (**Fig. 3B, Supplementary Fig. 6**). In contrast, constructs with C_1_-C_2_ distances of five and seven nucleotides led to a reduction in binding of at least one order of magnitude (**Fig. 3B**, **Supplementary Tab. 1**), confirming the findings of 6 nt as optimal C_1_-C_2_ distance.

### A computational algorithm to screen for the consensus in known She2p/3p substrates

We performed computational screening (details, see **Material and Methods**) to identify further instances of the SRS consensus motif in further target sequences of the She2p/3p transport machinery. For this purpose, we screened a list of 24 transcripts reported to show *in-vitro* binding by She2p/3p or bud localization *in-vivo*^24,29,30^. The identified sequence regions represented potential SRS instances and were evaluated for their potential to form the correct conformation of one of the configurations using *Kinefold*. We intended to extend this conformational evaluation by integrating data describing the degree of unpaired state of nucleotides (obtained in a remarkable genome-wide study using parallel analysis of RNA structure (PARS)^33^). This strategy was promising, but for our purposes the data were not detailed enough, due to competing conformations that could not be differentiated in the averaged values per nucleotide position, even for some of the strong *ASH1* SRS elements (**Material and Methods, Supplemental Fig. 7**).

For the three transcripts *EAR1, IST2* and *WSC2*, their SRS elements had been identified already, with their sequence regions narrowed down to fragments^10,30^. Now, with knowledge of the SRS consensus motif, these SRS elements could be classified as a config II arrangements (**Fig. 4C, Supplementary Fig. 8**). Remarkably, for *EAR1* and *IST2*, the asymmetric bulge was even more eccentric than observed in *ASH1* (*EAR1:* 7 nt: 0 nt; *IST2*: 9 nt: 1 nt). In addition, we identified an additional upstream SRS instance in each transcript, *EAR1* and *IST2* (**Tab. 2**). In contrast, for *WCS2*-C, the initially identified SRS fragment turned out to comprise possibly two SRS elements, a canonical one in the center of this fragment, and a potential SRS instance in the flanking regions, however, with a C_1_-C_2_ distance of 7 nt (**Supplemental Fig. 8D**). However, we could not classify the second published instance in *WSC2*, *WSC2*-N, in agreement with our model (see **Discussion**). As a first outcome of the screening we had identified seven SRS instances (**Fig. 3C**). When we studied the seven canonical SRS elements in detail, we noticed a recurring di-adenosine in the L-bulge (“AA_L_”) as a common feature in all elements (**Fig. 3C**), possibly indicating relevance in SRS function. This was surprising due to the fact that the crystal structure of E3 showed the interaction between She2p/3p and the AA_L_ nucleotides occurs exclusively via the backbone atoms of the nucleotides, and thus unspecifically^8^, with their nucleobases having intramolecular contact within the RNA structure. This observation makes relevance in the context of SRS recognition less likely.

**Figure 4.**
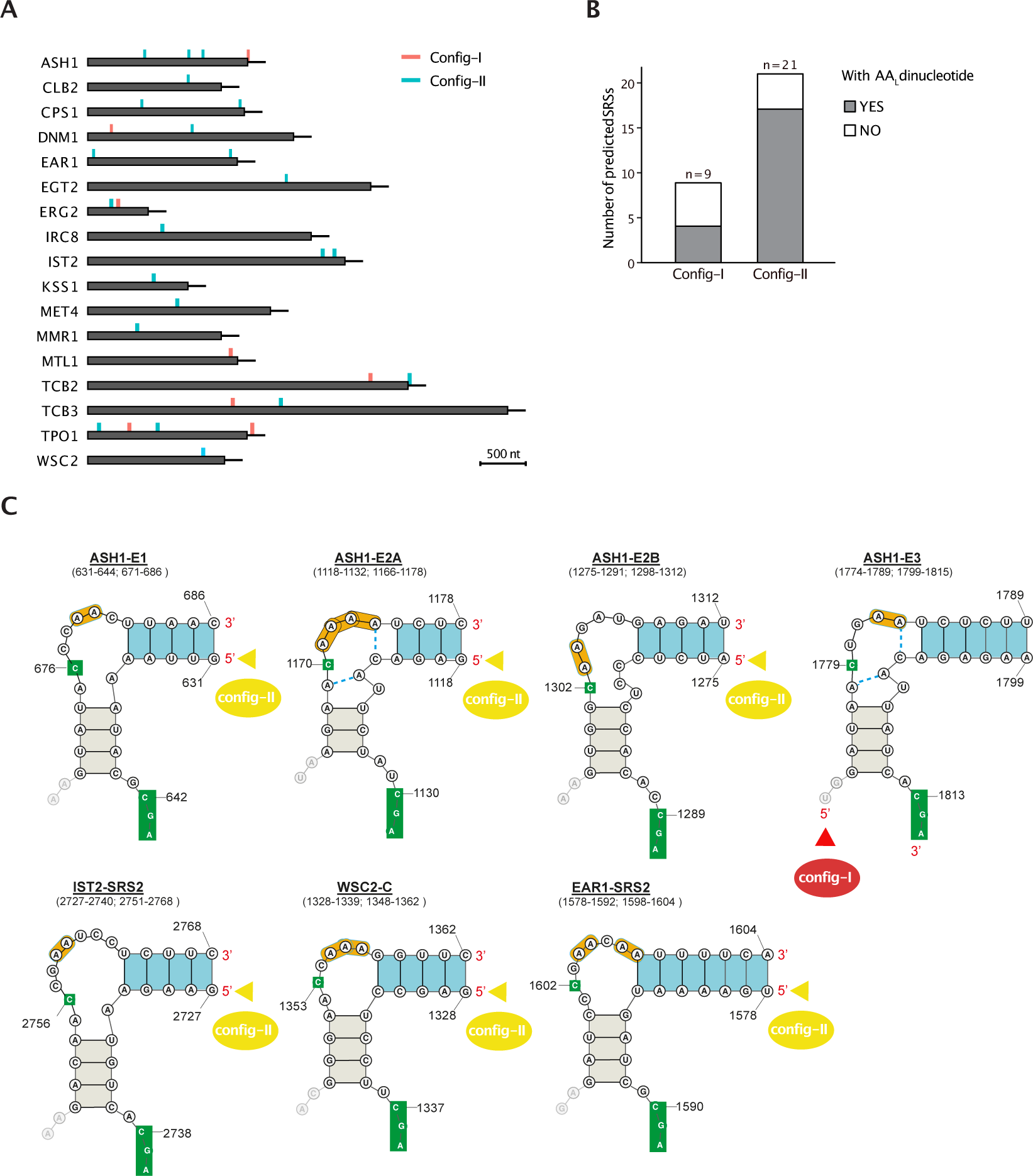
Computational screen for the SRS consensus motif across 24 known She2p/3p interacting transcripts results in 30 predicted instances in 17 transcripts. **A** Computational screen utilized both, feature screening of the SRS consensus motif and conformational evaluation (**Materials and Methods**) and was applied on 24 sequences known for She2p/3p interaction but with varying experimental confidence (**Supplementary Tab. 1**). It identified 30 SRS predictions in 17 sequences, here visualized as bar charts, with their coding sequences (rectangles) and 3’ UTR extensions (200 nt, black lines). SRS predictions were indicated by their configurations (config-I: red, config-II: blue). **B** For each configuration, the occurrence of the AA_L_ dinucleotide was specified. **C** Schematic overview on the high-confidential set of SRS elements. Similar to the SRS elements in *ASH1*, predictions in *IST2*, *EAR1* and *WSC2* matched published, well-characterized SRS elements (nomenclature, see **Tab. 2**) and could be arranged accordant to the model. Color scheme is adopted of Figure 1, with visualization of AA_L_ dinucleotides, boxed in orange). Uncommon base pairings had detected in the E3-She2p/She3p crystal structure were consistently mapped to E2A that displayed high sequence similarity with E3 after rotation (blue-dashed). Of note, similar uncommon base-pairings may also arise in E1 and *IST2*-SRS-2.

**TABLE 2.**
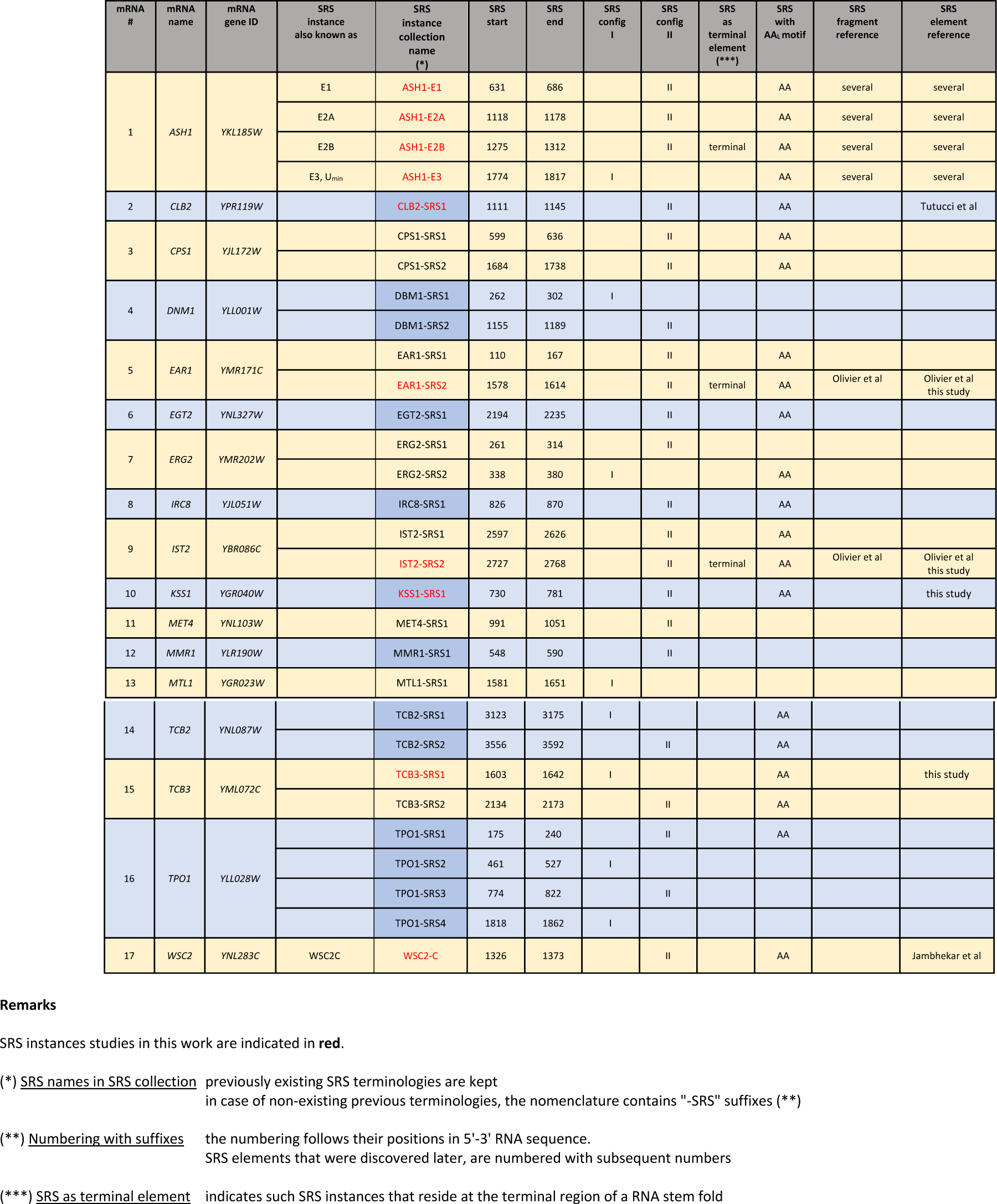
Overview on the *SRS instance collection.* The columns list info at transcripts and SRS instance level; SRS instance are classified according to the SRS consensus motif (including positions and configuration); the last two columns contain information to the identification of SRS elements in fragment size or as a detailed description in previous publications and this study

| mRNA # | mRNA name | mRNA gene ID | SRS instance also known as | SRS instance collection name (*) | SRS start | SRS end | SRS config I | SRS config II | SRS as terminal element (***) | SRS with AA motif | SRS fragment reference | SRS element reference |
| --- | --- | --- | --- | --- | --- | --- | --- | --- | --- | --- | --- | --- |
| 1 | ASH1 | YKL185W | E1 | ASH1-E1 | 631 | 686 |  | II |  | AA | several | several |
|  |  |  | E2A | ASH1-E2A | 1118 | 1178 |  | II |  | AA | several | several |
|  |  |  | E2B | ASH1-E2B | 1275 | 1312 |  | II | terminal | AA | several | several |
|  |  |  | E3, U<sub>min</sub> | ASH1-E3 | 1774 | 1817 | I |  |  | AA | several | several |
| 2 | CLB2 | YPR119W |  | CLB2-SRS1 | 1111 | 1145 |  | II |  | AA |  | Tutucci et al |
| 3 | CPS1 | YJL172W |  | CPS1-SRS1 | 599 | 636 |  | II |  | AA |  |  |
|  |  |  |  | CPS1-SRS2 | 1684 | 1738 |  | II |  | AA |  |  |
| 4 | DNM1 | YLL001W |  | DBM1-SRS1 | 262 | 302 | I |  |  |  |  |  |
|  |  |  |  | DBM1-SRS2 | 1155 | 1189 |  | II |  |  |  |  |
| 5 | EAR1 | YMR171C |  | EAR1-SRS1 | 110 | 167 |  | II |  | AA |  |  |
|  |  |  |  | EAR1-SRS2 | 1578 | 1614 |  | II | terminal | AA | Olivier et al | Olivier et al this study |
| 6 | EGT2 | YNL327W |  | EGT2-SRS1 | 2194 | 2235 |  | II |  | AA |  |  |
| 7 | ERG2 | YMR202W |  | ERG2-SRS1 | 261 | 314 |  | II |  |  |  |  |
|  |  |  |  | ERG2-SRS2 | 338 | 380 | I |  |  | AA |  |  |
| 8 | IRC8 | YJL051W |  | IRC8-SRS1 | 826 | 870 |  | II |  | AA |  |  |
| 9 | IST2 | YBR086C |  | IST2-SRS1 | 2597 | 2626 |  | II |  | AA |  |  |
|  |  |  |  | IST2-SRS2 | 2727 | 2768 |  | II | terminal | AA | Olivier et al | Olivier et al this study |
| 10 | KSS1 | YGR040W |  | KSS1-SRS1 | 730 | 781 |  | II |  | AA |  | this study |
| 11 | MET4 | YNL103W |  | MET4-SRS1 | 991 | 1051 |  | II |  |  |  |  |
| 12 | MMR1 | YLR190W |  | MMR1-SRS1 | 548 | 590 |  | II |  |  |  |  |
| 13 | MTL1 | YGR023W |  | MTL1-SRS1 | 1581 | 1651 | I |  |  |  |  |  |
| 14 | TCB2 | YNL087W |  | TCB2-SRS1 | 3123 | 3175 | I |  |  | AA |  |  |
|  |  |  |  | TCB2-SRS2 | 3556 | 3592 |  | II |  | AA |  |  |
| 15 | TCB3 | YML072C |  | TCB3-SRS1 | 1603 | 1642 | I |  |  | AA |  | this study |
|  |  |  |  | TCB3-SRS2 | 2134 | 2173 |  | II |  | AA |  |  |
| 16 | TPO1 | YLL028W |  | TPO1-SRS1 | 175 | 240 |  | II |  | AA |  |  |
|  |  |  |  | TPO1-SRS2 | 461 | 527 | I |  |  |  |  |  |
|  |  |  |  | TPO1-SRS3 | 774 | 822 |  | II |  |  |  |  |
|  |  |  |  | TPO1-SRS4 | 1818 | 1862 | I |  |  |  |  |  |
| 17 | WSC2 | YNL283C | WSC2C | WSC2-C | 1326 | 1373 |  | II |  | AA |  | Jambhekar et al |

As mentioned before, for most known SRS instances (in 23 of 30 instances in our set), functionality had been exclusively studied for large RNA fragments or complete transcripts. For 13 of these transcripts, we could identify now the positions of SRS instances. For 12 transcripts, we found one or two SRS instances in each sequence, and we found four instances in *TPO1* (**Fig. 4A**). Altogether, in 17 of 24 sequences (*ASH1*, *WSC2*, *EAR1*, *IST2* included), we predicted 30 SRS instances, with nine instances (30%) in config-I and 21 instances (70%) in config-II, respectively (**Fig. 4A**). Of notice, more than two thirds of the SRS instances contained the AA_L_ di-nucleotide in the Lbulge (**Fig. 4B**).

### Validation of SRS elements in *EAR1*, *IST2*, *KSS1* and *TCB3*

As described before, in *IST2* and *EAR1*, sequence fragments with SRS function had been identified already, and these sequences were also identified in our computational screening (termed *EAR1*-SRS2, *IST2*-SRS2; nomenclature see **Tab. 2**). We could classify their functional features, in agreement with the SRS consensus motif, provided that the consensus was valid for these instances. In such a view, the SRS elements would resemble the very first characterization (Olivier et al., 2005), however in the larger context of the SRS consensus motif. To address this aspect, we studied specifically designed SRS constructs, analogous to our previous studies. In accordance with the consensus, the newly designed constructs *EAR1_26nt_*switch and *IST2*_29nt_switch were strictly reduced to those regions assumed to be essential for SRS function and also switched in configuration (each to config-I) (**Fig. 5A**). As we could show in EMSA studies, *EAR1*_26nt_switch was bound with affinities comparable to the unmodified wild-type element *EAR1*_50nt_wt (**Fig. 5A**). Although for *IST2*_29nt_switch binding was clearly retained, it was considerably weaker when compared to *IST2*_62nt_wt. In addition, we tested the predicted *KSS1*-SRS1 (config-II, **Tab. 2**) and *TCB3*-SRS1 (config-I) for which no detailed position in their transcripts had previously been determined. As a result, we observed strong binding *in-vitro* for both, *KSS1*_72nt_wt and for the minimal construct KSS1_29nt_switch (switched to config-I), confirming *KSS1*-SRS1 as a potential SRS element (**Fig. 5B**). In an analogous manner, we observed for the minimal construct TCB3_38nt_wt (config-I) strong interaction with the fusion protein suggesting *TCB3*-SRS1 as further SRS element (**Fig. 5B**). In summary, we were able to demonstrate the identification and *in-vitro* confirmation of new SRS instances. Furthermore, for some of these instances we were able to prove that the newly designed, size-reduced constructs retain the SRS function even in switched configuration. In conclusion, the fact that our model, derived exclusively from *ASH1* SRS elements, enabled the identification, size reduction, and config-switch of designed SRS constructs in previously unexplored sequences while preserving SRS function confirmed its applicability as an SRS consensus motif.

**Figure 5.**
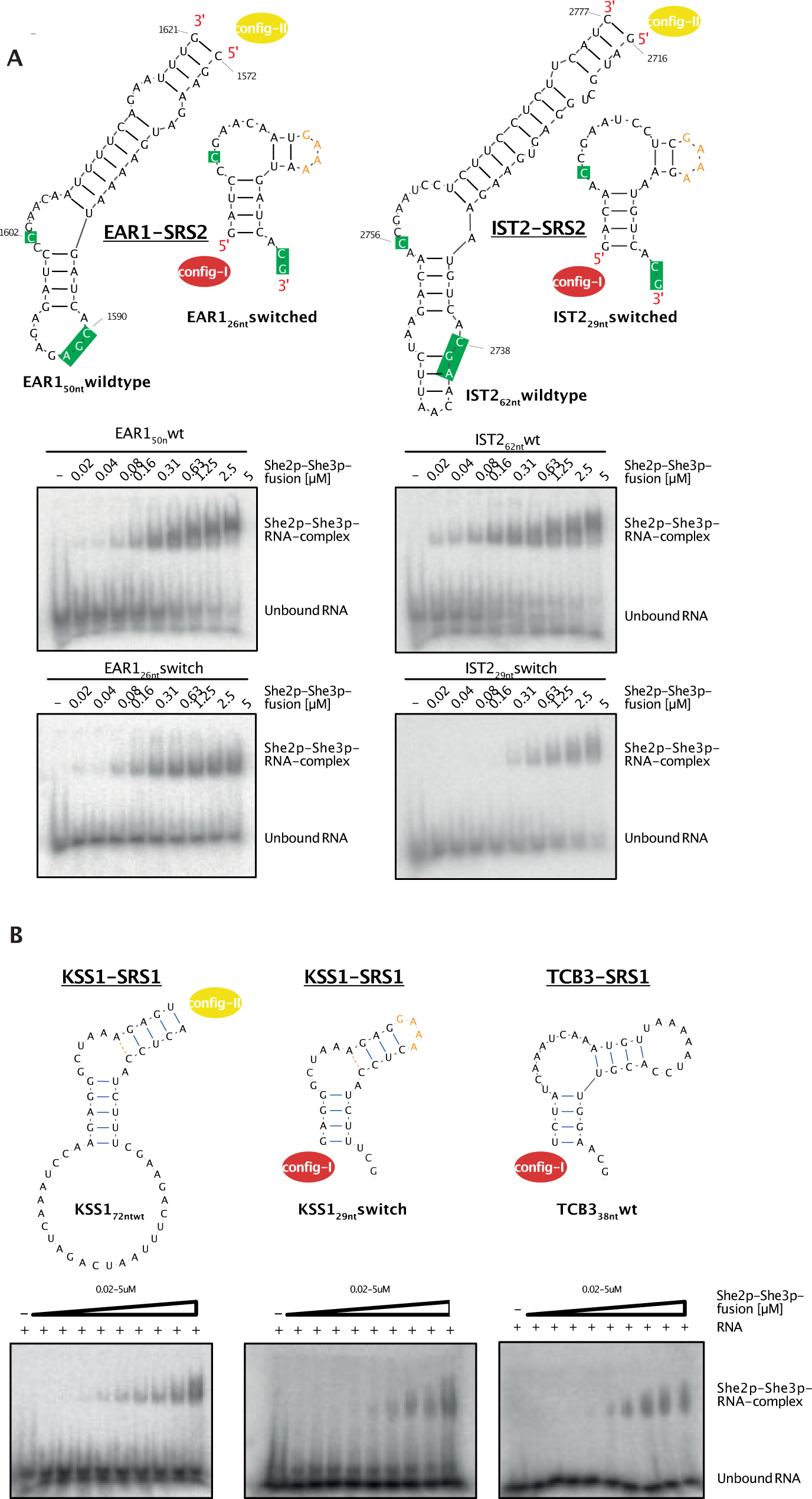
The configuration switch in the minimal SRS constructs of *EAR*-SRS2, *IST*-SRS2, *KSS1*-SRS1 and *TCB3*-SRS1 displays conserved SRS function in electrophoretic mobility shift assays. Representative results of electrophoretic mobility shift assays (EMSA) to analyze interaction of radioactive wild-type and designed RNA constructs with the She2p-She3p fusion protein (She2p_6-246_-(GGSGG)_2_-She3p_331-405_) in increasing concentration (0.02 - 5 μM). All minimal constructs were designed switched in configuration (from config-II to config-I). Their final loop sequence GAAA referred to the artificial loop sequence of the E3 constructs in ^8^ that was shown not to modify SRS function. **A** EMSA experiments of well-known SRS elements. Left, *EAR1*-SRS2_1572-1621_ (config-II) was studied as wildtype construct *EAR1*-SRS2_50nt_wt and as config-switched, minimal construct *EAR1*-SRS2_26nt-_switch (config-I). Right, *IST2*-SRS2_2716-2777_ (config-II) was studied as wildtype construct *IST2*-SRS2_62nt_wt and as config-switched, minimal construct *IST2*-SRS2_29nt-_switch (config-I). **B** EMSA experiments of new, predicted SRS elements. Left, *KSS1*-SRS1 (config-II) was studied as wildtype construct *KSS1*-SRS1_72nt_wt and as config-switched, minimal construct *KSS1*-SRS1_29nt-_switch (config-I). Right, *TCB3*-SRS1 (config-II) was studied as config-switched, minimal construct *TCB3*-SRS1_38nt-_switch (config-I).

### Validation of further features in the SRS of *EAR1*

In *EAR1* (*YMR171C*), it had been shown for its SRS element (now termed *EAR1*-SRS2) that substitutions in the sequence motifs decreased function^10^. The substitution of both, C_1_ and C_2_ (updated nomenclatures) in the constructs “M1” and “M2”^10^ had shown severe effects *in-vivo*. To examine the SRS consensus motif and *EAR1*-SRS2 in more detail, we chose a fine-grained approach using the *EAR1*_31nt_switch construct (switch to config-II) as the base construct for further changes (**Fig. 6A**) and determined its binding affinity using SPR analysis as described before. As result, we obtained an affinity of the original construct *EAR1*_31nt_switch of K_D_ = 11 nM (**Fig. 6D, Supplementary Fig. 9**). Remarkably, compared to the E3 minimal element, the affinity with the She2p/3p fusion protein was decreased by one order of magnitude (E3_29nt_wt: K_D_ = 0.89 nM). When we modified *EAR1*_31nt_switch in the inverted repeat IR1, by inversions or additionally introduced base pairs, the modified constructs showed affinities in the range of the original construct suggesting no direct impact on SRS function (**Fig. 6D**). Also, when we analyzed the relevance of the di-nucleotide AA_L_ in the Lbulge by deletions of different size (**Fig. 6C**), we observed binding affinities in the order of magnitude of the basic construct (**Fig. 6D**). The analysis of the functional cytosines C_1_ and C_2_ was complex since two cytosines (C_1601_ and C_1602_) precluded an unambiguous assignment of the C_1_ position (**Fig. 6B**). The substitution of either cytosine (constructs C_1-1_U, C_1-2_U) resulted in a considerable binding decrease suggesting that both positions contribute to She2p/3p binding (**Fig. 6D**). Unexpectedly, the deletion of one of the C_1_ positions (ΔC_1_) or a simultaneous substitution of both cytosines (C_1-1_ U & C_1-2_ U) restored She2p/3p binding (see **Discussion**). In contrast, substituting C_1590_ in the three-nucleotide C_1590_GA motif (C_2_U) returned unequivocal results manifesting in a dramatic loss of binding. Altogether, these results confirmed that neither sequence composition of IR1 nor the AA_L_ motif in the Lbulge were relevant for She2p/3p binding *in vitro*. Alterations of the C_1_-C_2_ distance were tolerated in some SRS contexts but not others, with a strong bias for a C_1_-C_2_ distance of six nucleotides. We therefore kept a C_1_-C_2_ distance constraint of six nucleotides for subsequent computational SRS screening.

**Figure 6.**
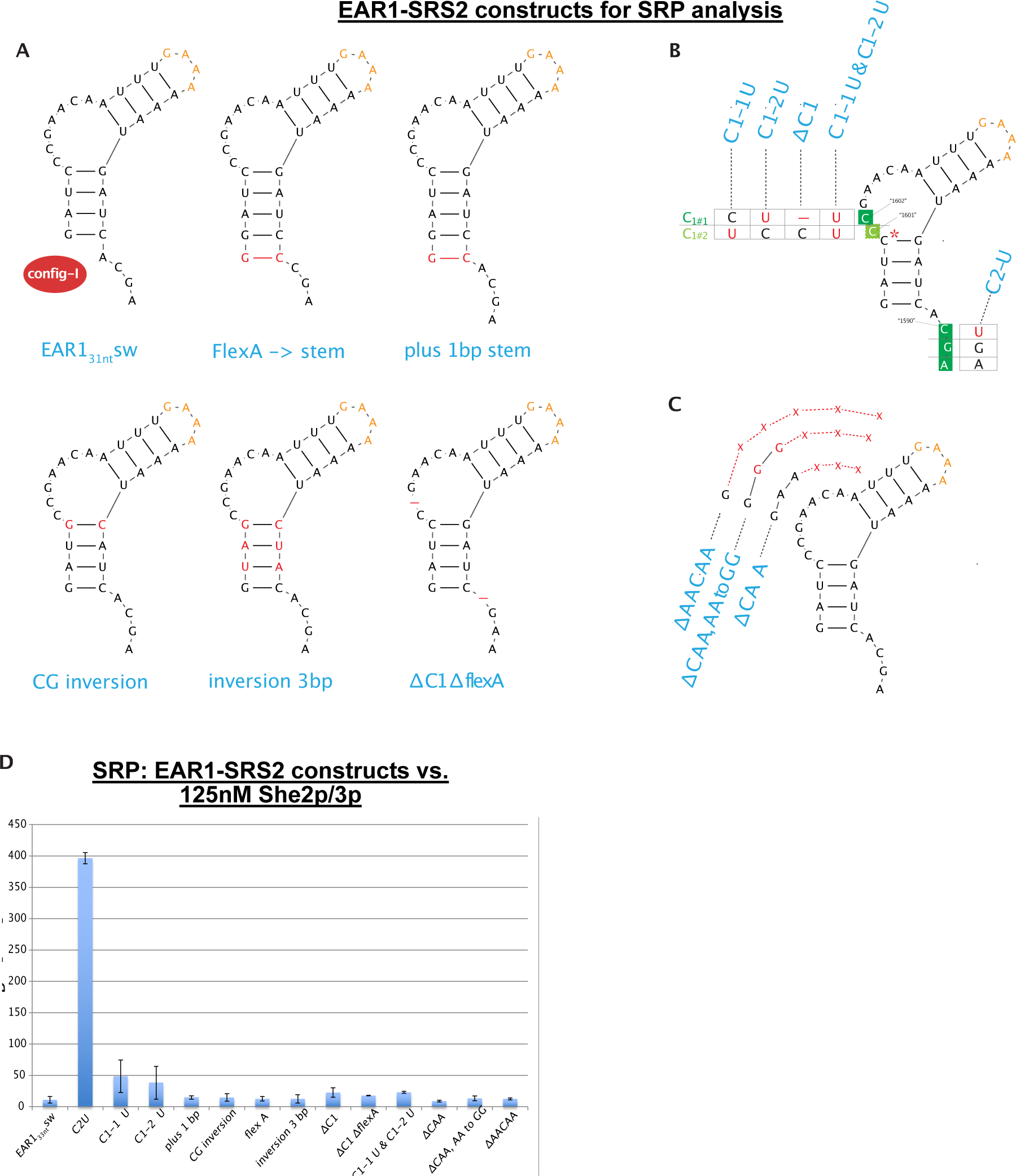
*EAR1*-SRS2_28nt_switch is partly sensitive to alterations of its functional motifs but tolerates alterations in its conformational features and in the AA_L_ dinucleotide of the Lbulge as showed by surface plasmon resonance analysis. A Schemes of the minimal constructs of *EAR1*-SRS2_28nt_switch that were re-designed to create constructs with altered conformational features (indicated in red in conformations). The final loop sequence GAAA referred to the artificial loop sequence of the E3 constructs in ^8^ that was shown not to alter SRS function **B** Substitution scheme to analyze the relevance of cytosines in the sequence motifs of the SRS consensus model. Alterations are marked in red in substitution tables that refer to both, sequence positions and terminologies of resulting constructs. Since two cytosines (C_1601_ and C_1602_) precluded an unambiguous assignment of the C_1_ position, both had to be considered. A third cytosine C_1600_, marked by a star, was not considered in alterations due to its uncommonly short distance to C_2_. **C** Substitution scheme to analyze the relevance of the AA_L_ dinucleotide in the Lbulge. For each alterations, construct name (blue), conserved sequence in the Lbulge (black) and alterations (red) are displayed. **D** Results of surface plasmon resonance analysis (SRP), performed in triplicates to analyze the minimal EAR1-SRS2 constructs and alterations in their C_1_-C_2_ distance. E3 RNA was incubated with She2p-She3p fusion protein (125 nM)

### A collection of predicted *SRS* instances in known target transcripts of She2p/3p

Computational screening for the SRS consensus motif had identified SRS instances in 17 of 24 sequences (**Fig. 4A**). After successful *in-vitro* validation of several of the new SRS instances (in *EAR1*, *IST2*, *KSS1*, *TCB3*), we finally compiled a collection of these screening results to present to the community (*SRS instance collection*, **Tab. 2**). This collection contains information on 30 SRS instances in 17 transcripts that have been reported to show She2p/3p binding *in-vitro* or bud localization *in-vivo* ^24,29,30^. In addition, a standardized nomenclature for SRS elements is introduced and further information is provided on positional data, classification and other details of SRS instances, several of them are promising new candidates for future experimental investigations.

## Discussion

In this work, we have derived the unified model of a three-dimensional RNA motif that functions as She2p/3p recognition motif (SRS) with the She2p/She3p protein transport machinery, a prerequisite for active transport of transcripts to the yeast bud. We used a computer-aided and conceptual modeling approach that based on the hypothesis that an asymmetric bulge in the RNA conformation might be a prerequisite of SRS function. This approach led to the identification of a common three-dimensional arrangement of SRS functional features (conformational features and sequence motifs) in the *ASH1* localization elements E3, E2A and E1 (and with limitations also for E2B) in a blueprint, thus demonstrating the long-missing consensus motif of all four SRS elements. Unexpectedly, although this arrangement of features was conserved in its overall three-dimensional fold, there were two variations with respect to the linear arrangement in the nucleotide sequences that we termed “configurations” (“config-I” in E3, “config-II” in E2A, E1 and E2B). The consensus model proved to be correct based on our consistent results from electric mobility shift assays, in which we analyzed the SRS function of RNA-constructs with re-designed feature arrangement. According to our model, this rearrangement of the features affected their linear order in the nucleotide sequences, but allowed the preservation of their overall three-dimensional arrangement, hence representing a switch between their original configuration and the respective counterpart configuration (“config-switch”). We were able to consistently demonstrate that the config-switch retained SRS function in all experiments we performed indicating that the consensus motif comprises the main requirements of SRS function. The fact that also size-reduced constructs proved to keep SRS function was a further hint that we had grasped the essential features of SRS function. It can be described in all *ASH1* SRS elements as follows: an asymmetric bulge, flanked by inverted repeats, forms a kinked, L-shaped RNA conformation. The larger side of the bulge contains the single cytosine motif C_1_ inside of the first three positions. The C_2_GA motif resides on the opposite strand of C_1_, separated by an inverted repeat in downstream direction on this strand. The two configurations are marked by the two different 5’ entry sites into this kinked conformation. These findings were further confirmed, when we compared previous thorough mutagenesis studies of the *ASH1* SRS elements and could assign these results well according to their relevance in the SRS consensus model. Further, by studying the binding kinetics of E3_29nt_wt in surface plasmon resonance analysis, we could confirm the optimal distance between C_1_ and C_2_ with six nucleotides (ca. 28 Å) as it was suggested before^10^. It is noteworthy that the existence of two configurations with double-stranded folds that can be converted into each other by swapping their features in the nucleotide sequence finds an analogy in studies on the beta-actin RNA zip code, where the order of its single-stranded 5’ and 3’ elements is interchangeable within the target transcripts bound to ZBP1^11^.

Due to its universal character to describe all four *ASH1* SRS elements, the exploitation of the SRS consensus motif went beyond previously computational screens. Hence, we created a computational screen that searched for both configurations, followed by a conformational check of the overall RNA fold performed by *Kinefold*. For some of the further transcripts that had been found to localize at the bud tip of yeast cells, sequence fragments had already been narrowed down with retained SRS function. In this set of sequence fragments, we could classify SRS elements of *EAR1*, *IST2* and *WSC2* as further instances of the SRS consensus motif. *EAR1* was first identified as localized in the bud in the study of ^29^ with its SRS element characterized in further studies^10^ and classified as a config-II SRS element in this work (*EAR1*-SRS2, **Tab. 2**). For the minimal construct *EAR1*_31nt_switch, using SRP analysis, we could confirm previous observations that the nucleotide sequences of the inverted repeats did not contribute to SRS function^8,13^. In further consistence with previous *in-vivo* studies^10^, the loss of C_2_ showed and individual substitutions of each of the two potential C_1_ positions (in C_1_-C_2_-distance of five or six nucleotides) affected SRS function. Unexpectedly, a single deletion of C_1601_ or a simultaneous substitution of C_1601_ and C_1602_, both restored SRS function. When studying the potential di-nucleotide AA_L_ sequence motif by deletion experiments we could not detect any impact on SRS function, even if the Lbulge region was almost completely deleted. So far, we can neither explain the apparent discrepancy in the C_1_ mutation effects in the *EAR1* constructs nor the retained SRS function with deleted asymmetric bulge. It may reflect both complex sequence composition and inherent redundancy in this particular SRS instance.

So far, we had observed only the occurrence of one config-I (E3) whereas we had identified six instances of config-II (**Fig. 4C**). This dominance of config-II SRS elements continued when we applied the computational screen to our complete sequence set and predicted 30 SRS instances in 17 transcripts, with 22 of them (73%) exhibiting config-II. So far, the reason for the unequal frequency of the two configurations remains unclear. From this outcome, we used EMSA approaches to analyze two instances of each subset: SRS instances with relatively well-known positions in their transcripts or with unknown SRS positions. For the first subset, we could confirm SRS function in both SRS elements *EAR1*-SRS2 and *IST2*-SRS2^10^. For the second subset, we could verify *KSS1*-SRS1 and *TCB3*-SRS1 *in-vitro.* Remarkably, when we switched configuration in *EAR1*-SRS2, *IST2*-SRS2 and *KSS1*-SRS1 (config-II to config-I), we could demonstrate again conserved SRS function for <u>all</u> switches, strongly supporting the SRS consensus model. Of note, as a first successful validation *in-vivo*, we had been able to identify and confirm the nucleotide positions of a SRS element in the *CLB2* transcript^34^: In the transcript, only a few but functionally crucial positions of the SRS instance *CLB2*-SRS1 (config-II, 1111 nt - 1145 nt) were altered in non-canonical mutations to reveal a significant loss-of-function in bud-localization. It is also interesting to note that the regulatory effect of well-defined asymmetric bulges seems not to be limited to SRS function. As described in a study^35^, the structure of the pre-mRNA autoregulatory complex of the L30 ribosomal protein in yeast displays an asymmetric bulge (length-ratio 5 nt: 2 nt) with a cytosine positioned in an extension of the Lbulge, and separated from another cytosine (in a further bulge) by an IR in a distance of 6 nt. Remarkably, it is this asymmetric bulge region that is part of a mutual induced fit, suggesting analogies with the observed induced fit mechanisms in the crystal structure of E3 and She2p/3p.

Further aspects of SRS characteristics remain to be investigated. For example, a further extension of the description of the SRS consensus motif is thinkable: It is reasonable to ask if the SRS function would be conserved if Lbulge and Sbulge were joined together without IR2 as intervening feature. Although experimental analysis of such a setting is beyond the scope of this study, we would like to put to discussion such a theoretical construct, to possibly take a step forward to complete the picture (**Supplementary Fig. 10A**). The SRS element *WSC2*-N^36^ that could be not classified, might be such an example (**Supplementary Figure 10B, left**). Such an extended perspective might have further implications also for the understanding of some of the SRP results of *EAR1* described before. Both elements, *EAR1*-SRS2 and *IST2*-SRS2 displayed <u>two</u> CGA motifs with the following consequences: Whereas we considered *EAR1*-SRS2 as config-II with C_2_GA residing in the terminal bulge, this terminal bulge also may be considered as a joint of Lbulge and Sbulge, hence offering a new perspective: the cytosine motif in this terminal bulge may be considered as C_1_ while its counterpart inside of the asymmetric bulge may be extended and labelled to C_2_GA (**Supplementary Figure 10B, right**). In such a view, the functionality of *EAR1*-SRS2 would exhibit redundant features of two superimposed and oppositely rotated SRS elements in the bigger context of an extended SRS consensus motif. This perspective may also explain partially the ambiguous results in the SPR experiments of our study. Secondly, a further extension of the description of the SRS consensus might be formed by considering a range in the restraints of the C_1_-C_2_ distance. In consistency with previous studies^10^ and with our SPR results we focused on the optimal parameters of six nucleotides in this study to minimize false positive matches. However, we could not exclude a minimal functionality for instances with (C_1_-C_2_ distance ≠ 6). An example might be a second, potential SRS instance that we identified in the flanking regions of *WSC2*-SRS2 (*WSC2*-C) but with a C_1_-C_2_ distance of seven nucleotides (**Supplementary Fig 7C**). In such a case, *WSC2*-SRS2 may compensate the suboptimal C_1_-C_2_ distance of its neighboring SRS element by inducing SRS conformation in a cooperative manner when both SRS instances interact with the binding interfaces on the opposite sites of the She2p tetramer^37^.

As a second aspect, localization elements that mediate bud localization require the interaction with the She2p/3p protein transport machinery, but further motifs may be required as binding platforms of auxiliary factors. For example, for E2B, we speculated that its energetically optimal conformation had to be converted to an alternative conformation to represent the SRS consensus motif. Even strong SRS elements such as E2A^12^ can be subject to major reorganizations, as shown by the PARS data which indicate competing conformations for E2A^33^ (**Supplementary Fig. 7**). Therefore, the She2p/3p interaction can be seen as a central step in the zip-code function, but it might be framed by further rearrangements in the chronological order. In such a scenario, transient binding of auxiliary cofactors could stabilize the asymmetric bulge and increase SRS function. AA_L_ might be such a co-factor binding motif. Despite the fact that AA_L_ was present in each case of our seven well-characterized, bud-localizing SRS elements, neither the crystal structure^8^ nor the results of SPR studies suggest involvement in She2p/3p binding specificity. Future research will be required to clarify the possible function of AA_L_ *in-vivo*, also with regard to transient interactions. In addition, for E3 and E2A, it was tempting to speculate if their extraordinary sequence identity was a product of convergent evolution, due to further requirements in affecting zip-code function. Maybe we have to face a trade-off between canonical SRS consensus motif and stability of the individual RNA conformation, with the weakening of one aspect may be compensated by strengthening of the other. This would complicate the identification of SRS instances, especially if auxiliary co-factors would alter the probabilities of formation of SRS conformations *in-vivo*. Summarizing, all these aspects stress two aspects: (i) The new SRS consensus motif delivers a conceptual framework of how to classify further aspects of new SRS instances, and (ii) It remains necessary to combine experimental screenings (identification of bud-localizing transcripts, final validation of instances) and computational screenings (prediction of positional information in transcripts to narrow down experimental analysis and suggest crucial positions in mutations studies). Consequently, the *SRS instance collection* contains only such SRS instances where bud localization had been confirmed already, at least at the transcript level.

In conclusion, the SRS consensus motif enabled the description of the four SRS elements of *ASH1* in one model, hence providing the long-missing consensus. It further unified the described functional features of the known SRS elements in *IST2*, *EAR1* and *WSC2* and enabled the identification of new SRS elements in further bud-localizing transcripts. The consensus model proved to be consistent with previous mutagenesis studies of these elements and we could demonstrate that a switch between the configurations retained function in SRS constructs of several transcripts.

## Materials and Methods

### Overview on the theoretical data flow to derive the SRS consensus model

The theoretical part of deriving the SRS consensus model comprised the following steps (**Supplementary Fig. 2**): (1) Hypothesis (asymmetric bulge as hallmark of SRS) by analysis of crystal structure of *ASH1*-E3 (2) Selection of an appropriate conformational prediction tool (3) Conformational prediction of instances and inspection to derive a consensus (**Fig. 2**) (4) Cross-check of the consensus with published functional data (5) Creation of a computational screening pipeline (6) Check for further data sources to filter results (**Supplementary Fig. 7**) (7) Screening for instances in selected sequence sets (8) Inspection of instances for further patterns

### Identification of an appropriate conformational prediction tool

The inspection of the crystal structure of *ASH1*-E3 suggested an asymmetric bulge as a pre-requisite of kinked RNA conformation. The occurrence of such asymmetric bulges may be favorized in mRNPs by the energy release due to molecular interaction. However, its occurrence is less likely in an individual RNA and is less likely to be predicted as plausible conformation by computational prediction tools that commonly deal with individual RNA sequences. In a survey, when investigating which of common tools^32,38–40^ was most likely to predict the asymmetric bulge of E3, we found Kinefold^32^ as the one that best tolerated asymmetric bulges (**Supplementary Fig. 3**). It predicted the asymmetric bulge consisting of a 5 nt Lbulge and a 2 nt Sbulge, which closely approximates the length ratio of the asymmetric bulge observed in the E3-She2p/3p crystal structure^8^ that displayed a length ratio of 6 nt: 3 nt (considering only Watson-Crick base pairs) or 4 nt: 1 nt (considering also atypical base pairings as observed in the crystal structure), respectively (**Fig. 1A**).

### Cross-check of the consensus with published functional data

We used published data of functional studies that involved both, mutagenesis and deletion of RNA regions to cross-check for consistency with the SRS consensus model^10,13,25,30^. We characterized such mutations in *ASH1* SRS elements that could clearly be assigned to destroy or to keep SRS function according to the consensus model. All these constructs were in accordance with the consensus model (data not shown).

### Development of a computational screening approach

For the purpose of identifying further instances of the SRS consensus motif, we transformed the consensus features into “patterns” to screen for “matches” in RNA sequences. We created a computational screening pipeline consisting of in-house computational scripts and an external conformational prediction algorithm. For the screening, three aspects were required: (1) Screening the sequence for conformational aspects. For each RNA, the sequence was screened for two nested pairs of inverted repeats (each with a minimal length of four nucleotides) that framed an asymmetric bulge region. We had to consider both configurations in different screening patterns. The obtained constituted potential SRS instances. (2) Screening the sequence for functional motifs. For each SRS instance, the Lbulge region was screened for C_1_, the opposite strand for C_2_GA (positioned beyond IR1), with a C_1-_C_2_ distance of 6 nt between. The screening strategy based on the approach of modular feature screening of mosaic-like patterns^41^. This modular approach allowed to introduce further features to be screened (e.g., AA_L_). It also tolerated length variations between the distinct (sequence and conformational) features (e.g., one version tolerated C_1_ positioned in the last nucleotide of IR1). As a consequence, permutated sets of varying patterns could be summarized into two screening patterns (one for each configuration). (3) Validation by conformational prediction. For each SRS instance, it was calculated if it would express a stable conformational fold that fitted to the conformational arrangement of the SRS consensus. This analysis was performed using *Kinefold* as energy-minimization approach that had proven best for large asymmetric bulges. On each site, the SRS stretches were extended by 10 nt of the flanking sequences of their RNA sequences, to tolerate some degree of freedom during calculation. In a series of tests using known SRS elements, a flanking region extension by 10 nt had proven to be best in a range of 0 nt to 50 nt (data not shown). Of note, natural limitations of common RNA conformational predictions occur due to the fact that such calculations are limited to predict the conformation of distinct RNA stretches but less reliable regarding statements of the complete transcript and second, they would ignore any energetic benefits by binding of molecular interactors (e.g., She2p/3p proteins). As a consequence, we analyzed the 25th best conformational predictions of *Kinefold* (maximum value = 25), to consider slight structural dynamics in each RNA molecule.

### Check for further data sources to filter for the consensus

For the purpose to possibly extend the validation step of new SRS instances by conformational predictions. we intended to cross-check the predicted conformations with experimental data to get deeper insights on actual RNA conformations *in-vivo*. For our purposes, we tested the remarkable genome-wide study based on PARS (Parallel Analysis of RNA Structure)^33^. For each nucleotide in transcripts, PARS-scores described the percentage in occurring in single-strand (“S1-reads” of RNase S1) or double-strand (“V1-reads” of RNase V1) regions of the RNA conformations. We extracted the S1- and V1-data and tested the strong localization element E3^13^ as positive control. As a result the PARS data was well suited to describe the SRS conformation as observed in the E3 crystal structure^8^ (**Supplemental Fig. 7**). However, for further controls, the data comparison of paired/ unpaired occurrence of nucleotides between PARS data and SRS consensus model failed (**Supplemental Fig. 7**). Commonly, RNA molecules display not one distinct conformation in-vivo but are subject to conformational dynamics resulting in collections of competing conformations. PARS data per nucleotide averaged the binding state (paired/unpaired) over all competing conformations. It seemed that even the two SRS elements in *ASH1*-E1 and *ASH1*-E2A (proven to be efficient localization elements^13^) were not sufficiently stable in the consensus conformations according to their PARS data to distinguish them from alternative conformations

### Further bioinformatics resources

For the investigation of molecular structures, we used Pymol^42^ as 3D-viewer. Initial protein characterization of She2p and She3p was performed using the ELM server^43^. RNA sequence alignments were performed using Jalview^44^. RNA sequences were extracted of the Saccharomyces Genome Database^45^. For the actual screening, the mRNA sequences of *ASH1*, *IST2*, *EAR1* (aka *YMR171C*), *WSC2* and 20 further mRNA sequences known to She2p/3p interact^10,13,25,29,30,36^ were chosen. As sequence data, we used the CDRs, extended by 200 nt downstream by default, due to the fact that for several mRNAs, the actual 3’ UTR lengths were still unknown. Functional annotation of Supplementary Table S1 followed ^46^. Graphical visualization of RNA conformations was performed using VARNA^47^.

### Protein expression

Expression in *E. coli* and purification of She2p, She3p, their fusion protein, and variants thereof were performed as described in ^8,23^. Purity and solubility were confirmed by size-exclusion chromatography and SDS page.

### RNA preparation and isotope labeling

For small scale *in vitro* transcription the MEGAshortscript T7 kit (life technologies) was used according to the manufacturer’s instructions. Synthetic DNA oligonucleotides containing a T7 promoter and the respective sequence were used as a template. After *in vitro* transcription, the template was digested by DNAse and the RNA was extracted using phenol:chloroform:isoamyl alcohol (25:24:1). Homogeneity of RNA was assessed by native and denaturing TBE agarose gel electrophoresis.

### Electrophoretic mobility shift assay

In a total volume of 20 µL 5 nM radioactively labeled RNA, 100 µg/mL yeast tRNA competitor, and protein at a given concentration were mixed in buffer (20 mM Hepes, pH 7.8, 200 mM NaCl, 2 mM MgCl_2_, 2 mM DTT) with 4 % (v/v) glycerol. After 25 min at room temperature, the samples were resolved by native TBE PAGE (6 % polyacrylamide) and gels incubated for 15 min in fixing solution (10 % (v/v) acetic acid, 30 % (v/v) methanol) and subsequently vacuum dried. Gels were analyzed with radiograph films or by a using Phosphorimager (FujifilmFLA5100).

### Surface plasmon resonance analysis (SPR)

SPR experiments were performed with a BIACORE 3000 system (GE Healthcare). For protein-RNA interactions, biotinylated RNA was streptavidin-captured on an SA-Chip (GE Healthcare) surface according to manufacturers instructions. Measurements were performed at a flow rate of 30 µL/min in standard buffer. Data were analyzed using the BIAevaluation software (GE Healthcare). Sample sensograms were referenced against the signal in a ligand-free reference channel, and averaged values at the equilibrium calculated. Steady state binding curves were obtained by plotting the response units at equilibrium against protein concentration with the Origin Pro 9.0 (OriginLab) software.

### Computational quality check of the folds of the experimental constructs

Part of the design of experimental constructs (minimal, config-switch) for the *in-vitro* and *in-vivo* approaches was a quality check with *Kinefold*: The conformations of their sequences were predicted to ensure that their expected conformation would be in agreement with the SRS consensus model

## Acknowledgements

We would like to thank Herve Isambert for helpful discussions regarding Kinefold and Toby Gibson for further support. This work was supported by the Deutsche Forschungsgemeinschaft (DFG, German Research Foundation) to R.-P.J. (DFG-FOR2333, Project Number 270067186), to M.F. (CEPLAS EXC 1028, CRC1208, DFG-FOR2333, DFG Fe448/8, DFG Fe448/9), to D.N. (DFG-FOR2333, Project Number 270067186), and to Z.K. (DFG-FOR2333, Project Number 270067186).

## INFO-BOX Summary of the SRS consensus motif

1. Two in**verted repeats** (IR-1, IR-2) flank an **asymmetric bulge** (consisting of **Lbulge** and **Sbulge**)
2. The conserved **singular cytosine C_1_** resides in one of the first three positions of the Lbulge
3. The conserved **C_2_GA triplet** resides in a further bulge, at the opposite strand of C_1_, separated by an inverted repeat in downstream direction on this strand in a **C_1_-C_2_ distance of six positions**
4. The two **configurations** are marked by the two different 5’ entry sites into the kinked SRS element conformation.

