## supplemental for "A consensus motif in *ASH1* and further transcripts unifies several RNA motifs required for interaction with the She2p/She3p transport machinery and mRNA localization in yeast"

[#] Equal contribution

[*] Corresponding authors

### Supplementary Figures

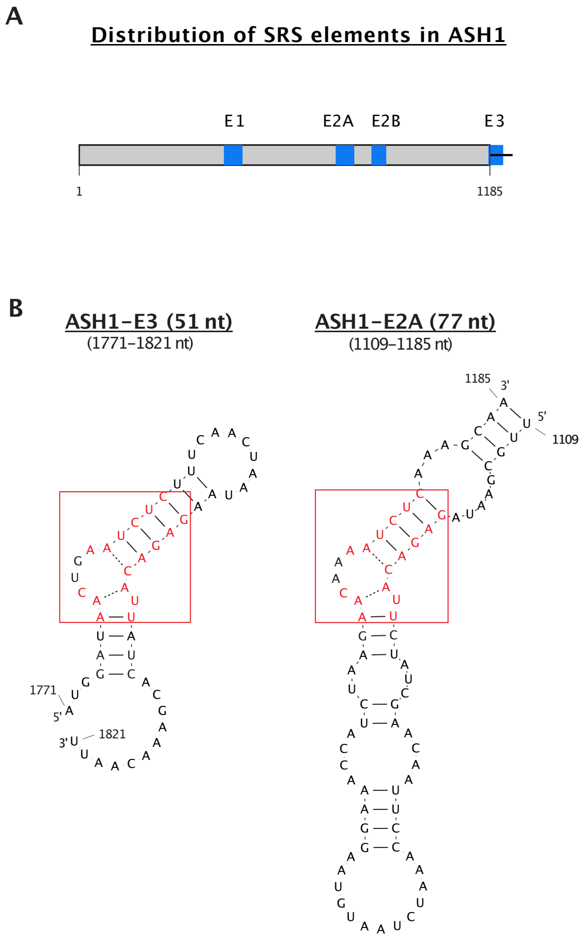


**Supplementary Figure S1. Distribution of the four localization elements in *ASH1* and sequence identities in E3 and E2A. A** In the sequence of the *ASH1* transcript, the three localization elements E1, E2A and E2B are positioned within the coding region, whereas E3 overlapped the stop codon and extended into the 3' UTR. **B** After adaption of the conformational predictions of the two SRS elements E3 and E2A by 180 degree rotation, the SRS elements display remarkable sequence identies (with fragments boxed in red, degree of sequence identity: 87%).

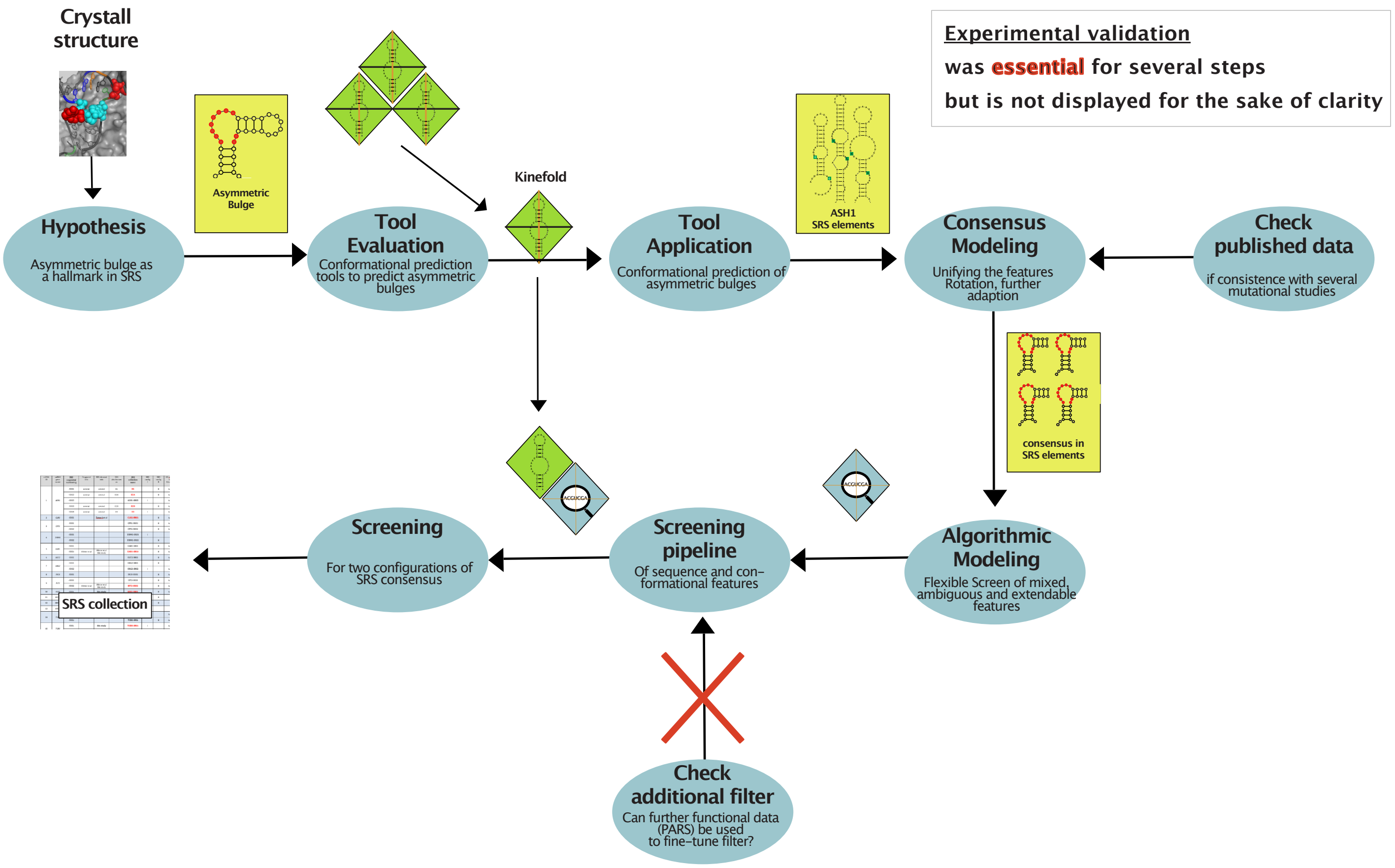


**Supplementary Figure S2. The theoretical concept of data flow towards the SRS consensus motif and the SRS instance collection.** Starting point has been the crystal structure[1], showing the SRS element E3 in interaction with She2p/She3p and leading to the hypothesis of the central asymmetric bulge of E3 as a requirement of SRS function. Several conformational prediction tools were evaluated to test which one may predict such asymmetric bulges in the best way, even for single RNA sequences. Using the conformational prediction tool *Kinefold* (displayed as green diamond) optimized the prediction of respective asymmetric bulges in the four *ASH1* SRS elements and could be used to model the SRS consensus motif finally. The cross-check between previously published mutational studies of the four *ASH1* SRS elements and the SRS consensus motif proved its consistence. The features of this model (sequence motifs, conformational features, configurations) were integrated in a computational screen (displayed as magnifier icon), preconnected by an overall conformational check by Kinefold. The test if additional external data may be feasible to be used as further filters was not successful. Using the computational screen allowed to identify further potential SRS instances in transcripts known to exhibit SRS function that were integrated into the "SRS instance collection".

### Validation of conformational prediction tools

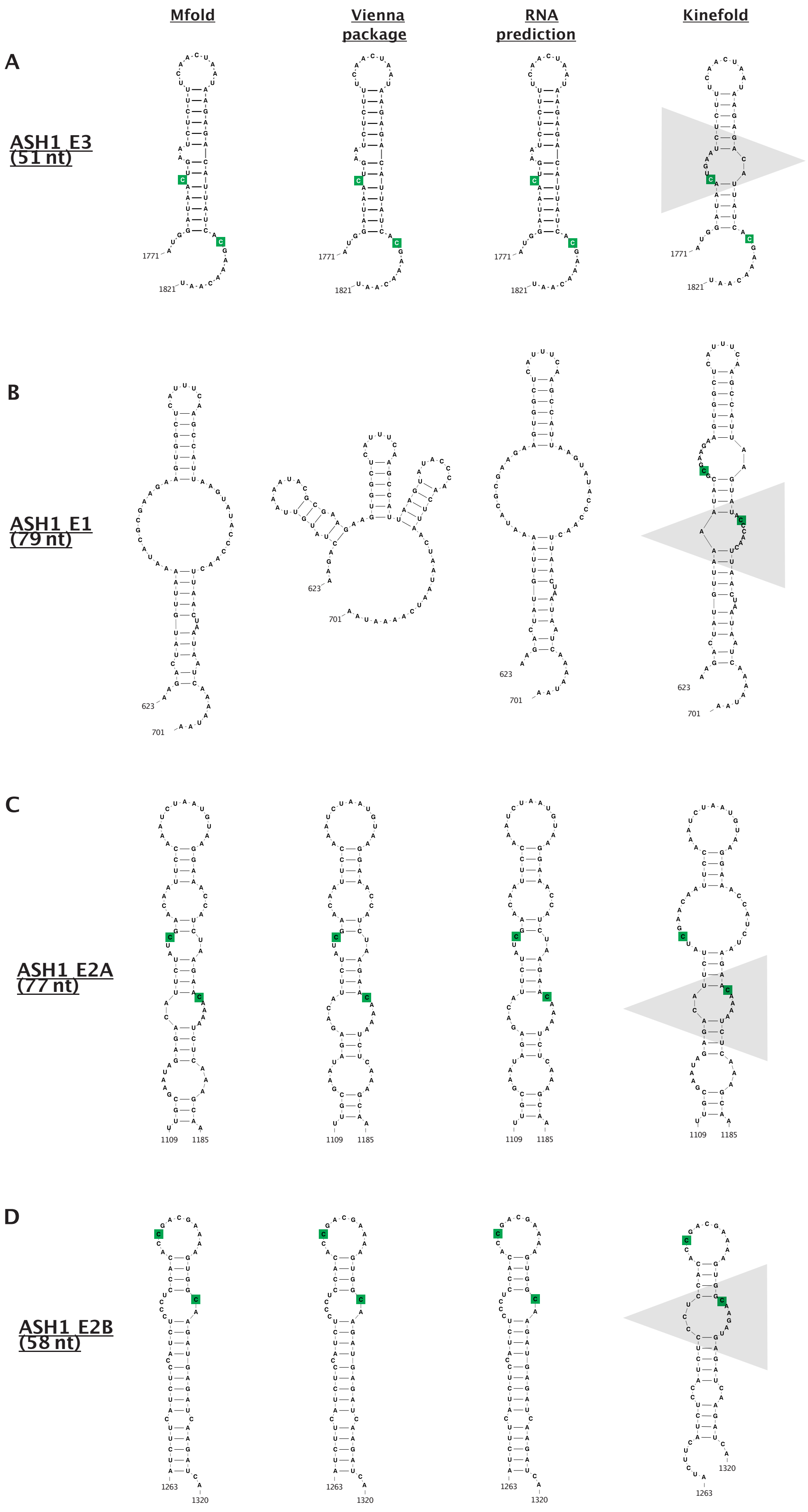

**Supplementary Figure S3. Validation of RNA conformational prediction tools.** The applications Mfold[2], Vienna package[3], RNA prediction[4] and Kinefold[5] were tested for their potential to predict asymmetric bulges in the four *ASH1* SRS elements. Finally, it was Kinefold that proved to fit best in the prediction of such conformations that displayed asymmetric bulge that also contained known functional cytosine motifs - as observed for E3 in the E3-She2p/3p crystal structure before. For the Kinefold predictions, the respective asymmetric bulges are indicated by a wedge colored in grey.

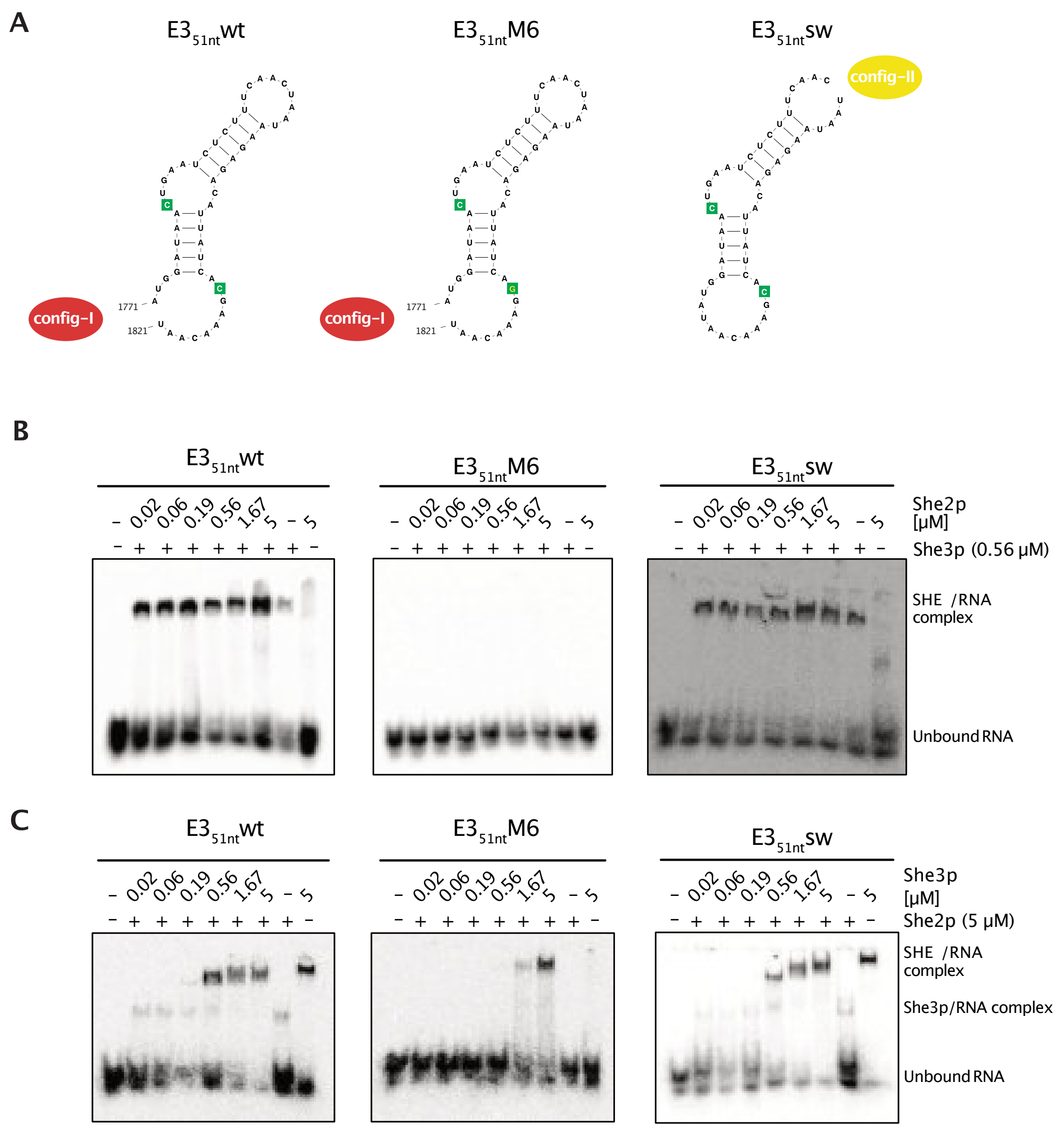


**Supplementary Figure S4. The configuration switch in E3 displays conserved function in electrophoretic mobility shift assays by incubation with full-length She2p and She3p proteins. A** Overview on conformational folds of the E3 constructs used in electrophoretic mobility shift assays (EMSA). **B** Representative results of EMSA triplicates to analyze complex formation of wild-type and variant constructs of radioactive ASH1 E3 51nt RNA with constant She3p concentration of 0.56 µM and titration of She2p (0.02 - 5 µM). **C** Representative results of EMSA triplicates as shown in Supplementary Figure S2-B, but with constant She2p concentration (5 µM) and titration of She3p (0.02 - 5 µM).

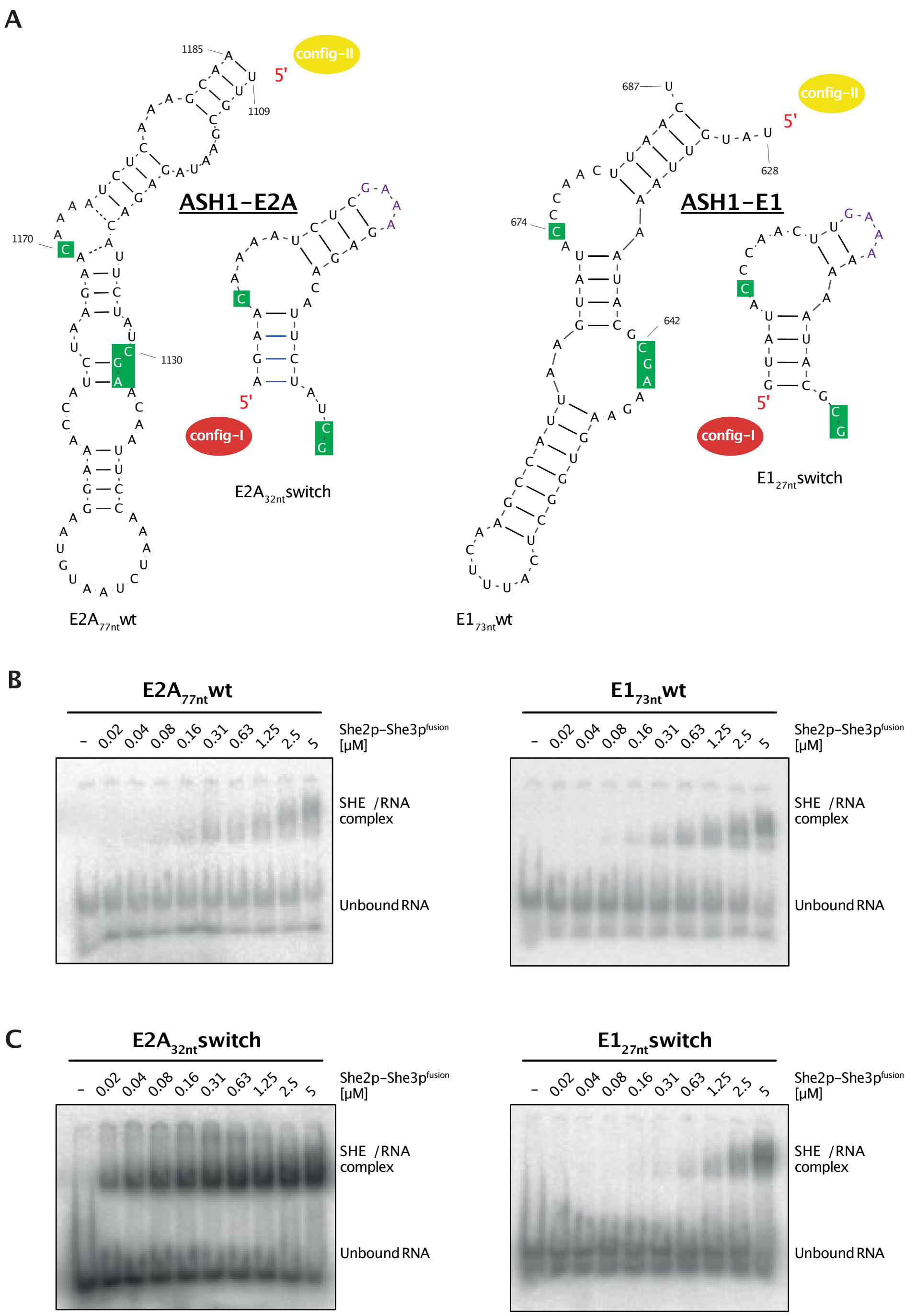


**Supplementary Figure S5. E2A and E1 displays conserved SRS function even for the config-switch. A** View on the conformation of the examined SRS constructs in their wildtype configuration (config-II) $E2A_{77nt}$wt and $E1_{73nt}$wt and in their switched configuration (config-I) $E2A_{32nt}$switch and $E1_{27nt}$switch with the functional cytosine motifs indicated in green . **B** Representative results of electrophoretic mobility shift assays (EMSA) of triplicates to analyze complex formation of the wild-type constructs with the She2p-She3p fusion protein (She2p6-246-(GGSGG)2-She3p331-405) in increasing concentration (0.02 - 5 µM). **C** Representative results of EMSA triplicates to analyze complex formation of the config-switch constructs; conditions as indicated in B.

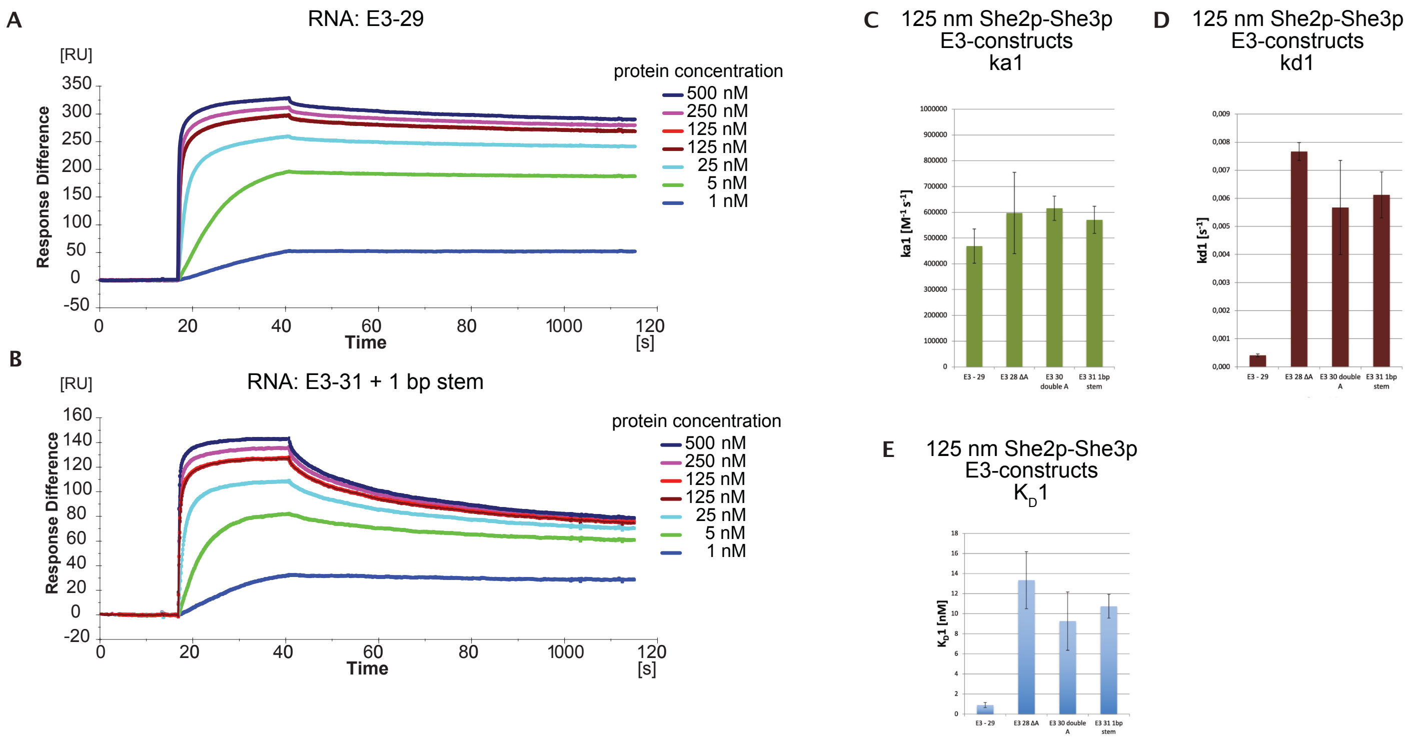


**Supplementary Figure S6. Surface plasmon resonance analysis confirms the optimal $C_1$-$C_2$ distance of 6 nt in minimal constructs of E3.** Results of surface plasmon resonance analysis (SRP), performed in triplicates to analyze interaction with She2p/3p fusion proteins with the minimal E3 constructs and alterations in their $C_1$-$C_2$ distance. E3 constructs refer to the conformation views of Fig. 3B. **A** Exemplarily graph of the outcome of the E3-29 nt wildtype sequence after incubation with She2p-She3p fusion protein in different concentrations (1nM to 500 nM). **B** Exemplarily graph of the outcome of the mutated construct E3-31 nt (one additional 1bp stem included) after incubation with She2p-She3p fusion protein in different concentrations (1nM to 500 nM). **C, D** $k_a1$ and $k_d$ 1 values of the different E3 constructs, incubated with 125 nM She2p-She3p fusion protein, were derived by bivalent analyte fit. **E** Display of the resulting KD1 values resulting of $k_a1$ and $k_d$ 1 values of C and D.

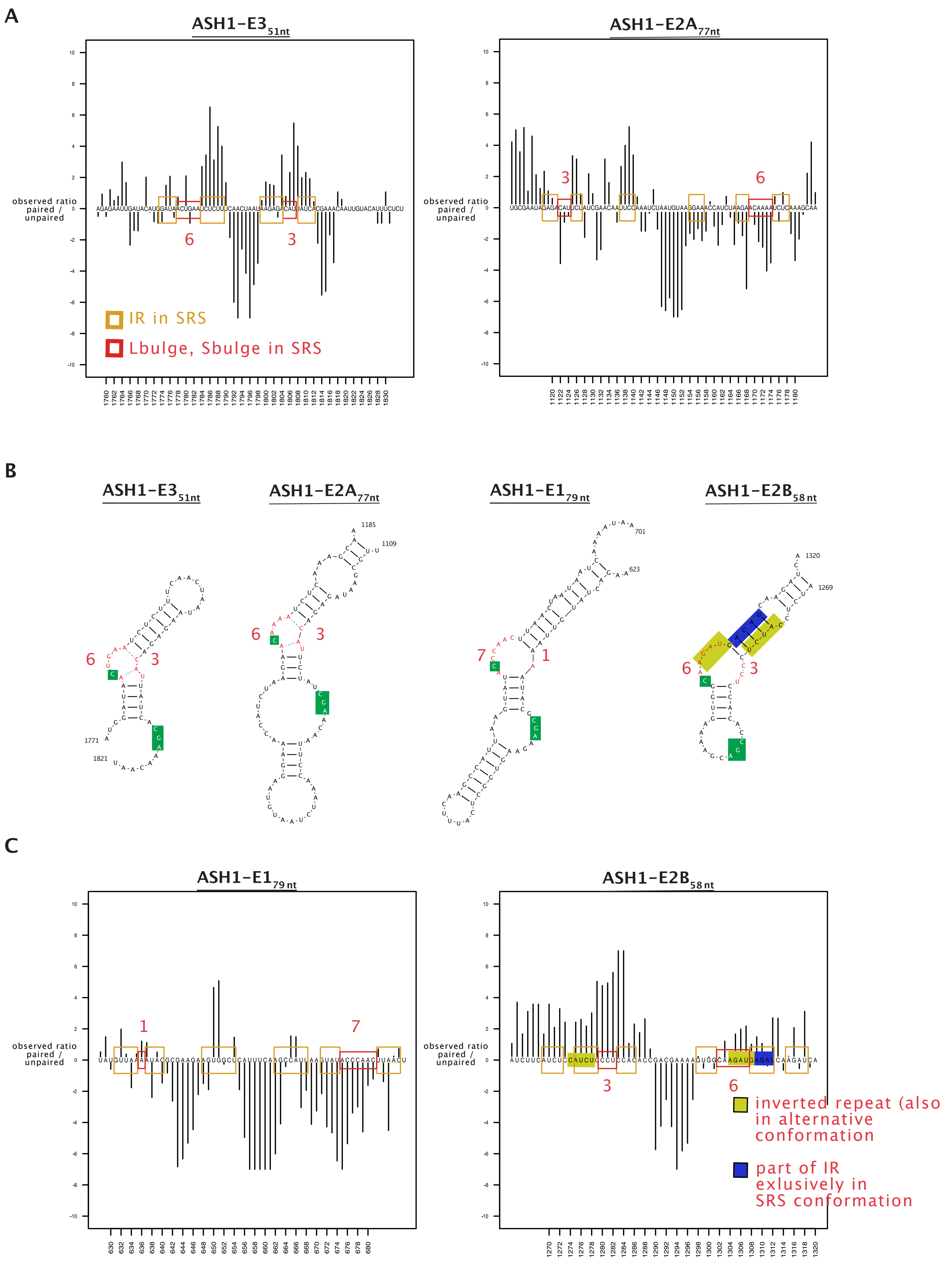
A
ASH1-E3 51nt
ASH1-E2A 77nt
observed ratio paired / unpaired
IR in SRS
Lbulge, Sbulge in SRS
B
ASH1-E3 51nt
ASH1-E2A 77nt
ASH1-E1 79 nt
ASH1-E2B 58 nt
C
ASH1-E1 79 nt
ASH1-E2B 58 nt
observed ratio paired / unpaired
inverted repeat (also in alternative conformation
part of IR exlusively in SRS conformation

**Supplementary Figure S7. Further external data are not suitable as further filtering criteria in the computational screen due to competing conformations that can not be differentiated in the averaged values per nucleotide position.** For the purpose to add a further filtering step in our computational screening pipeline we tested PARS information (Parallel Analysis of RNA Structure)[6], available at a transcriptome-wide scale. PARS-scores of each nucleotide position were graphically displayed, with positive values (referring to the baseline) indicating nucleotides in double-stranded structures. **A** Display of PARS scores of the *ASH1* SRS localisation elements E3 and E2A. The positions of inverted repeats are boxed in orange according to the SRS consensus motif (conformations displayed **B**). The positions of Lbulge and Sbulge are boxed in red and their lengths are indicated in red numbers. **C** Display of PARS scores of the *ASH1* SRS localisation elements E1 and E2B. Regarding E2B we have postulated, that the SRS conformation would have to evolve from an alternative conformation by the shifting of inverted repeats. For this setting, sequences of the alternative IR are indicated in yellow (in **B, C**) while the new sequence being involved in the IR of the SRS conformation is indicated in blue.

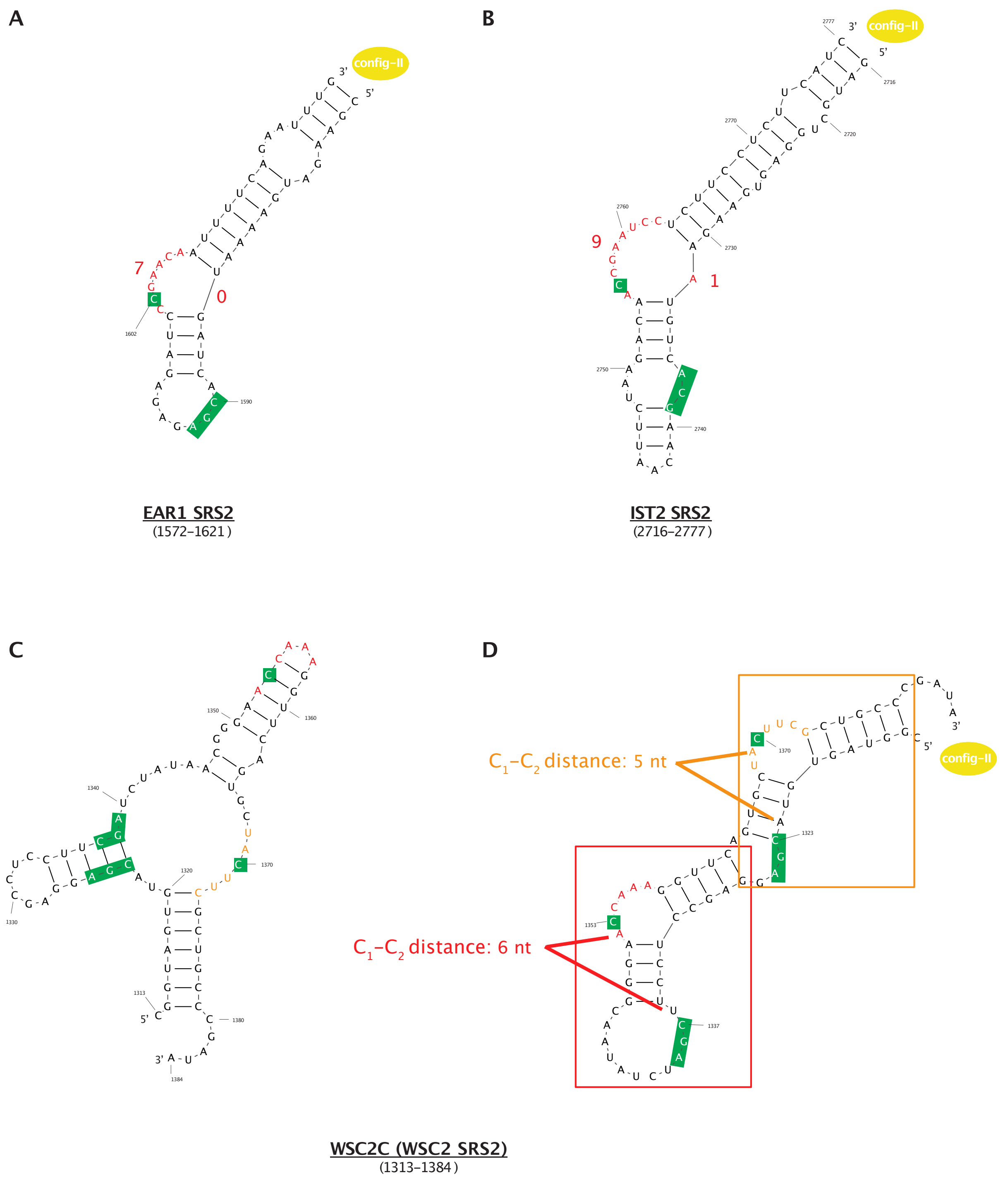


**Supplementary Figure S8. Classification of the SRS elements EAR1 SRS2, IST2 SRS2 and WSC2 SRS2 according to the SRS consensus motif.** The potential cytosine sequence motifs are marked in green, the Lbulge regions of the asymmetric bulges are marked in red. **A** Conformational view on SRS element EAR1 SRS2. **B** Conformational view on SRS element IST2 SRS2. **C** Conformational view on SRS element WSC2C as published in[7]. Functional sequence elements are indicated referring to D. **D** Conformational view on SRS element WSC2-SRS2 that displays a potential second SRS instance in its flanking regions. The conformation displays a canonical SRS element according to the SRS consensus model (red boxed) as well as a second, potential non-canonical SRS element, with a $C_1$-$C_2$ distance of 5 nucleotides in the respective flanking regions (orange boxed). Both (potential) SRS elements display a config-II arrangement.

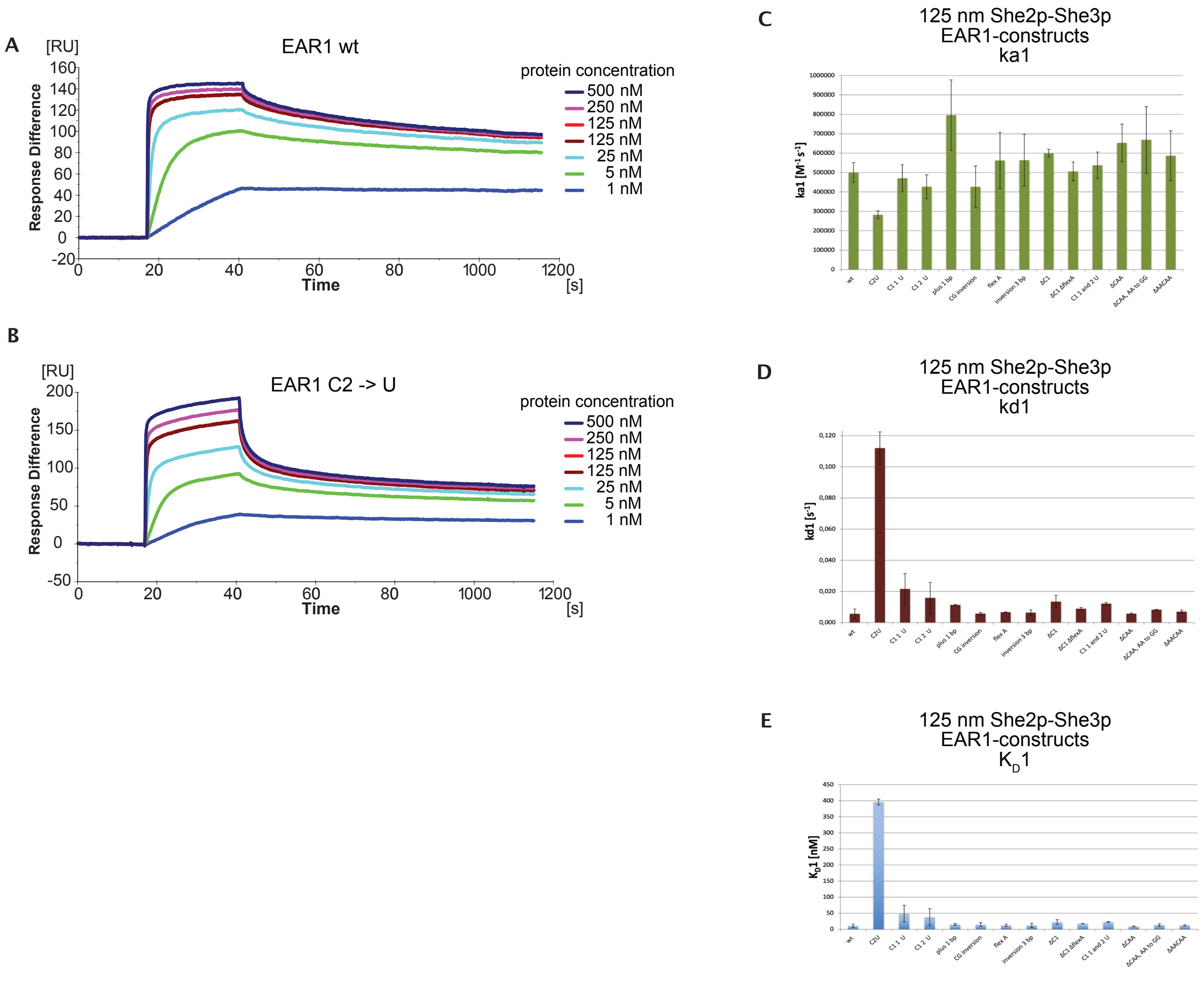


**Supplementary Figure S9.** ***EAR1*-SRS2 is partly sensitive to alterations of its functional motifs but tolerates alterations in its conformational features and in the AAL dinucleotide of the Lbulge as confirmed for its minimal, config-switched construct by surface plasmon resonance analysis.** Results of surface plasmon resonance analysis (SRP), performed in triplicates to analyze interaction with She2p/3p fusion proteins with the minimal *EAR1*-SRS constructs and alterations. *EAR1*-SRS2 constructs refer to the conformation views of Fig. 6A-C. **A** Exemplarily graph of the outcome of the *EAR1*-SRS2 wildtype sequence after incubation with She2p-She3p fusion protein in different concentrations (1nM to 500 nM). **B** Exemplarily graph of the outcome of the mutated construct *EAR1*-SRS2 C2>U after incubation with She2p-She3p fusion protein in different concentrations (1nM to 500 nM). **C, D** $k_a1$ and $k_d1$ values of the different *EAR1*-SRS2 constructs, incubated with 125 nM She2p-She3p fusion protein, were derived by bivalent analyte fit. **E** Display of the resulting KD1 values resulting of $k_a1$ and $k_d1$ values of C and D.

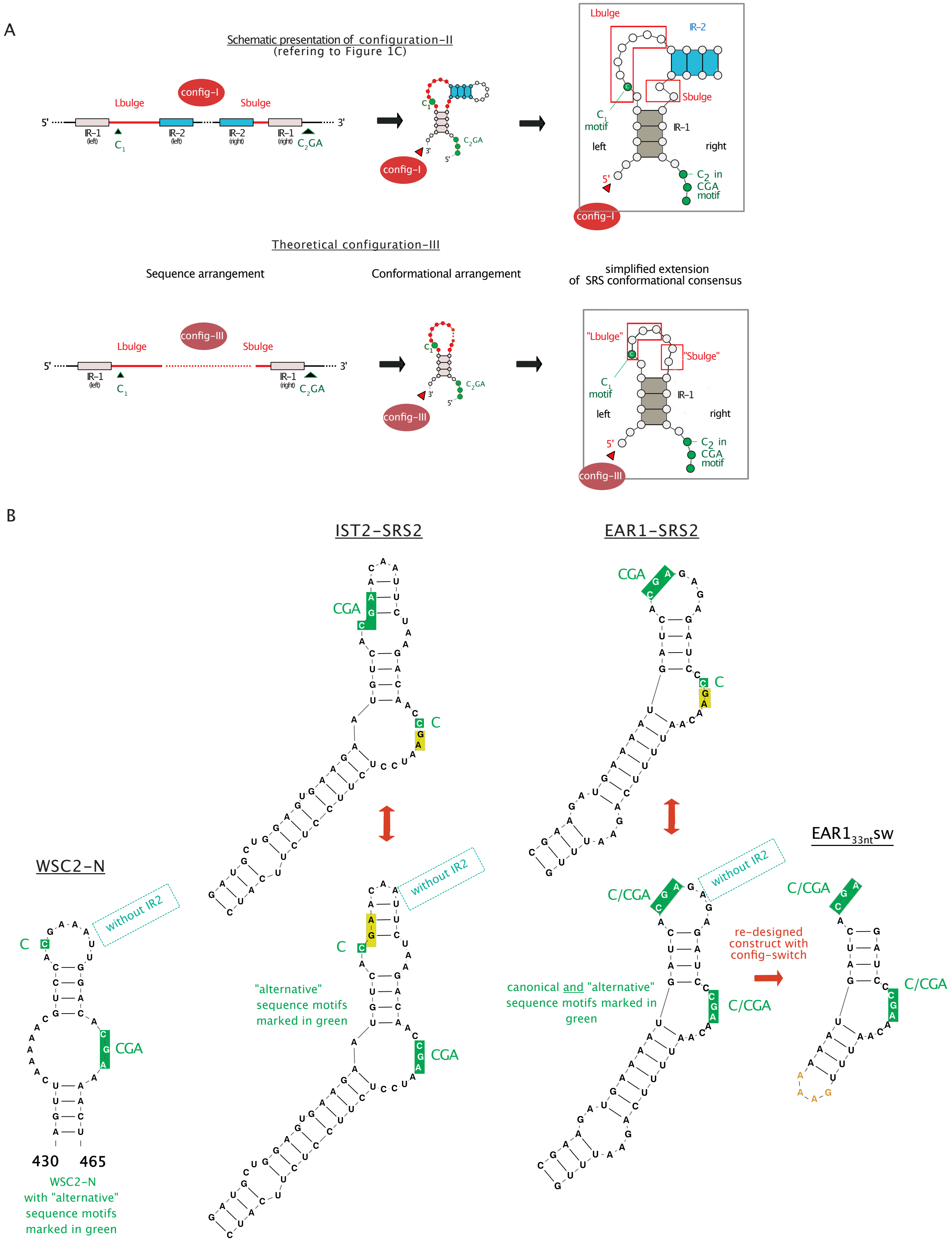
A
Schematic presentation of configuration-II
(refering to Figure 1C)
config-I
Lbulge
Sbulge
5'
3'
IR-1 (left)
IR-2 (left)
IR-2 (right)
IR-1 (right)
C1
C2GA
Lbulge
IR-2
Sbulge
C1 motif
IR-1
left
right
C2 in CGA motif
Theoretical configuration-III
Sequence arrangement
Conformational arrangement
simplified extension
of SRS conformational consensus
config-III
"Lbulge"
"Sbulge"
B
IST2-SRS2
EAR1-SRS2
WSC2-N
EAR1 33nt SW
CGA
C
without IR2
C/CGA
"alternative"
sequence motifs
marked in green
canonical and "alternative"
sequence motifs marked in
green
re-designed
construct with
config-switch
430
465
WSC2-N
with "alternative"
sequence motifs
marked in green

**Supplementary Figure S10. Schematic presentation of the SRS consensus model and how a potential extension of the SRS consensus model, derived of config-II may be feasible. A** Top row displays a schematic presentation of the SRS consensus model with its features displayed in config-I (IR1 boxed in grey, IR2 boxed in blue, Lbulge and Sbulge in red, sequence motifs in green, referring to Fig.1C). Bottom row displays a potential extension of config-II, derived by omitting inverted-repeat-2. As a consequence, in its conformational view, this potential config-III displays all consensus features besides of IR-2. **B** Conformational view of SRS instances displaying the potential config-III: function of the identified SRS element WSC2N[7] may be interpreted by the presence of a config-III arrangement. For both SRS elements, *EAR1*-SRS2 and *IST2*-SRS2, it is feasible to overlay onto the features of the config-II arrangement a second config-III arrangement, supposing config-III would be functional without the respective, missing inverted repeats-2. Such a config-III may also be understood as a config-I arrangement with missing IR-1 (as indicated in the figure in blue). In such a view, some of the minimal EAR1-SRS2 constructs would still be functional, despite of mutations in the config-II arrangement, due to redundant config-III features.

**Supplementary Tables**

**Supplementary Table S1. Compilation list of transcripts reported to interact with the She2/3p transport machinery**. Reference data indicate experimental work that identified transcripts as targets of She2p/She3p

| | Name | Gene name | Length CDS | Function | Reference 1 Takizawa 2000 [8] | Reference 2 Shepard 2003 [9] | Reference 3 Jambhekar 2005 [7] | Number predicted SRS elements |
|---|---|---|---|---|---|---|---|---|
| 1 | ASH1 | YKL185W | 1767 | Transcriptional regulator | Y | Y | Y | 4 |
| 2 | BRO1 | YPL048W | 2535 | Vacuolar protein sorting factor | | Y | | 0 |
| 3 | CLB2 | YPR119W | 1476 | B-type cyclin | | Y | | 1 |
| 4 | CPS1 | YJL172W | 1731 | Vacuolar Carboxypeptidase | | Y | | 2 |
| 5 | DNM1 | YLL001W | 2274 | Dynamin-related GTPase | | Y | Y | 2 |
| 6 | EAR1 | YMR171C | 1653 | Rsp5p-dependent ubiquitination | | Y | | 2 |
| 7 | EGT2 | YNL327W | 3126 | Cell wall endoglucanase | | Y | | 1 |
| 8 | ERG2 | YMR202W | 669 | C8-sterol isomerase | Y | Y | Y | 2 |
| 9 | IRC8 | YJL051W | 2469 | Unknown function | Y | Y | | 1 |
| 10 | IST2 | YBR086C | 2841 | Tethering of ER plasmamembrane | Y | Y | | 2 |
| 11 | KSS1 | YGR040W | 1107 | Mitogen-activ. protein kinase | | Y | | 1 |
| 12 | LCB1 | YMR296C | 1677 | Sphingolipid synthesis | | Y | | 0 |
| 13 | MET4 | YNL103W | 2019 | Transcription factor | | Y | | 1 |
| 14 | MID2 | YLR332W | 1131 | Sensor cell wall integrity | | Y | | 0 |
| 15 | MMR1 | YLR190W | 1476 | Myo2 adaptor | Y | Y | | 1 |
| 16 | MTL1 | MTL1 | 1656 | Sensor cell wall integrity | | | | 1 |
| 17 | PDR3 | YBL005W | 2931 | Transcription activator | Y | Y | | 0 |
| 18 | RGL1 | RGL1 | 1440 | Regulator of Rho1 | | Y | | 0 |
| 19 | SRL1 | YOR247W | 633 | Cell wall Mannoprotein | Y | Y | Y | 0 |

| | | | | | | | | |
|---|---|---|---|---|---|---|---|---|
| 20 | TAM41 | YGR046W | 1158 | Phosph Cytid Transferase | Y | Y | | 0 |
| 21 | TCB2 | YNL087W | 3537 | Tethering of ER plasmamembrane | | Y | | 2 |
| 22 | TCB3 | YML072C | 4638 | Tethering of ER plasmamembrane | | Y | | 2 |
| 23 | TPO1 | YLL028W | 1761 | Polyamine transporter | Y | Y | Y | 4 |
| 24 | WSC2 | YNL283C | 1512 | Stress sensor transducer | Y | Y | Y | 2 |
